# Super-enhancers shape the landscape and repair dynamics of transcription-associated DNA breaks in cancer

**DOI:** 10.1101/2025.05.25.655982

**Authors:** Osama Hidmi, Diala Shatleh, Sara Oster Flayshman, Jonathan Monin, Rami I. Aqeilan

## Abstract

Cancer is characterized by uncontrolled proliferation accompanied by the hypertranscription of oncogenes, leading to transcription stress, a key source of DNA double-strand breaks (DSBs) that jeopardize genomic stability. Yet, transcription stress is still underexplored. In this study, we utilized maps of DSBs identified through in-suspension break labeling *in situ* and sequencing (sBLISS), along with transcription stress markers, revealing that transcription stress regions coincide with the super-enhancer regulatory landscape. Notably, γH2AX mapping indicates its enrichment at transcription stress sites, while not all DSB-enriched genes show equal γH2AX marking, but those with DSBs tied to transcription stress are distinctly marked. Intriguingly, genes with high-DSBs marked by γH2AX exhibited significantly higher DSB turnover and repair than those with γH2AX-low genes, manifesting vulnerability to mutagenesis. These findings underscore super-enhancer activity as a determinant of the transcription stress landscape in cancer, posing a threat to the genomic stability of oncogenes.

## Introduction

The most fundamental hallmark of cancer cells is their ability to sustain proliferation, which they acquire through genetic alterations during neoplastic transformation^1^. Simultaneously, dysregulation of the transcriptional program emerges to accommodate the cell’s increased demand for cellular factors essential for proliferation, growth, and survival^2,3^. Consequently, genes encoding these factors, i.e., oncogenes, are hypertranscribed to fulfill the needs of cancer cells^4^.

A well-established mechanism by which cancer cells upregulate oncogenes involves the formation of oncogenic super-enhancers (SEs)^5–12^. In normal cells, SEs drive the expression of cell-type-specific genes that encode master transcription factors (MTFs), which shape cellular identity and the transcriptional program^2^. In cancer cells, SEs drive the expression of cancer-type-specific oncogenes and oncogenic MTFs that dictate their transcriptional programs and support their proliferation^2,12^. SEs can become dysregulated in cancer through various mechanisms: 1) overexpression of enhancer transcription factors renders SEs much more effective at driving the expression of their associated genes; 2) translocations can relocate SEs near an inactive oncogene in the corresponding normal cell, and 3) focal amplifications of SEs can lead to the increased expression of target genes^12^. These mechanisms contribute to the activation of proto-oncogenes by SEs, which drive the transcription of these genes to an exceptionally high level termed “hypertranscription.”

Transcription is increasingly viewed as a source of DNA damage^13–18^. The most damaging type of DNA lesions produced during transcription is DNA double-strand breaks (DSBs)^19^. These DSBs often result indirectly from the processing of transcription-associated intermediates, such as single-strand breaks (SSBs) that form during the repair of trapped topoisomerase 1 cleavage complexes (TOP1cc). DSBs resulting from transcription are primarily facilitated by the formation of pathogenic co-transcriptional R-loops and the aberrant activity of topoisomerase 1 (TOP1)^13,20^. R-loops tend to form more frequently at highly expressed genes, and their formation causes TOP1 to become trapped in its cleavage complex form (TOP1cc)^13^.

Although transcription is a highly regulated process, it can occasionally become abortive and result in DSBs. These transcriptional aberrations occur more frequently as transcription levels increase, in a phenomenon known as “transcription stress”^21^. Characterizing the “hotspots” of transcriptional stress is crucial because transcription-related DSBs can lead to cell death, genomic instability, and mutations^22,23^. Moreover, since cancer cells upregulate genes vital for their proliferation and survival, a deeper understanding of transcriptional stress and transcription-associated DSBs associated with these oncogenes can aid in exploiting their vulnerabilities to target cancer cells more effectively. Nevertheless, while transcription stress has been increasingly studied^24–26^, its landscape and consequences, particularly in cancer, remain incompletely characterized.

DSBs are impediments to transcription^27,28^ and must, therefore, be efficiently repaired to sustain continuous high transcriptional levels. However, it is not understood how regions of transcription stress are equipped to handle these deleterious DSBs.

In this study, we characterize the landscape of transcription stress in cancer cells and reveal how these regions are structured to promote the rapid and efficient repair of transcriptional DSBs that sustain hypertranscription.

## Results

### Super-enhancer-regulated oncogenes are major transcription stress sites

Pathogenic R-loops preferentially form at highly expressed genes^29^, facilitating aberrant TOP1 activity and TOP1cc trapping^13^. The high TOP1 occupancy, TOP1cc trapping, and co-transcriptional R-loops all contribute to the formation of transcription-associated DSBs ^20,30^. Therefore, we reasoned that these features, along with genome-wide maps of endogenous DSBs (hereafter referred to as “Breakome”), could serve as markers of transcriptional stress. To determine transcription stress sites in the genome, we analyzed sBLISS and ChIP-seq data for the breast cancer cell line MCF7 and applied a Hidden Markov Model (HMM)-based approach to identify regions in the genome with co-occurrence across the four datasets (sBLISS, DRIP-seq, TOP1cc, and TOP1 ChIP-seq). As expected, the majority of these regions were genic (Extended Data Fig. 1A), while the rest were enriched mainly at intergenic super-enhancers (SEs) (Extended Data Fig. 1B and C). To investigate which genes serve as transcription stress sites, we scored each gene based on TOP1, TOP1cc, R-loops, and DSBs scores (see Methods). This scoring yielded a shortlist of genes, many of which have well-established oncogenic functions (Fig. 1A). Upon closer analysis of this list, we noted that many of the genes were regulated by super-enhancers (SEs) in MCF7 cells, including CCND1, NEAT1, miR21, and MYC (SE-regulated genes were downloaded from SEdb 2.0^31^, see methods) (Fig. 1A). To statistically validate this observation, we systematically assessed the representation of multiple gene sets among transcription stress sites by performing gene set enrichment analysis (Fig 1B; see Methods). SE-regulated genes were the most significantly enriched category, with SE-regulated oncogenes exhibiting the highest enrichment (∼122-fold over expectation under a random model, P = 1.85 × 10^-15^; hypergeometric test). Similar results were obtained when quantifying the overlap between each gene set and the co-marked regions identified by HMM (Extended Data Fig. 1D-E). Notably, the enrichment of SE-regulated genes was substantially higher than that of typical-enhancer (TE)-regulated genes or highly expressed genes lacking enhancer regulation (Fig. 1B) despite having similar expression levels (Extended data Fig. 1F). To control for factors that might confound this analysis, we performed multivariate logistic regression using SE-status, Expression level, GC content, and gene length as predictors for transcription stress at genes. The results of this model demonstrate that SE-status was the most statistically significant independent predictor of transcription stress (Extended Data Fig. 1G), indicating specificity to genes regulated by SEs.

**Fig. 1:**
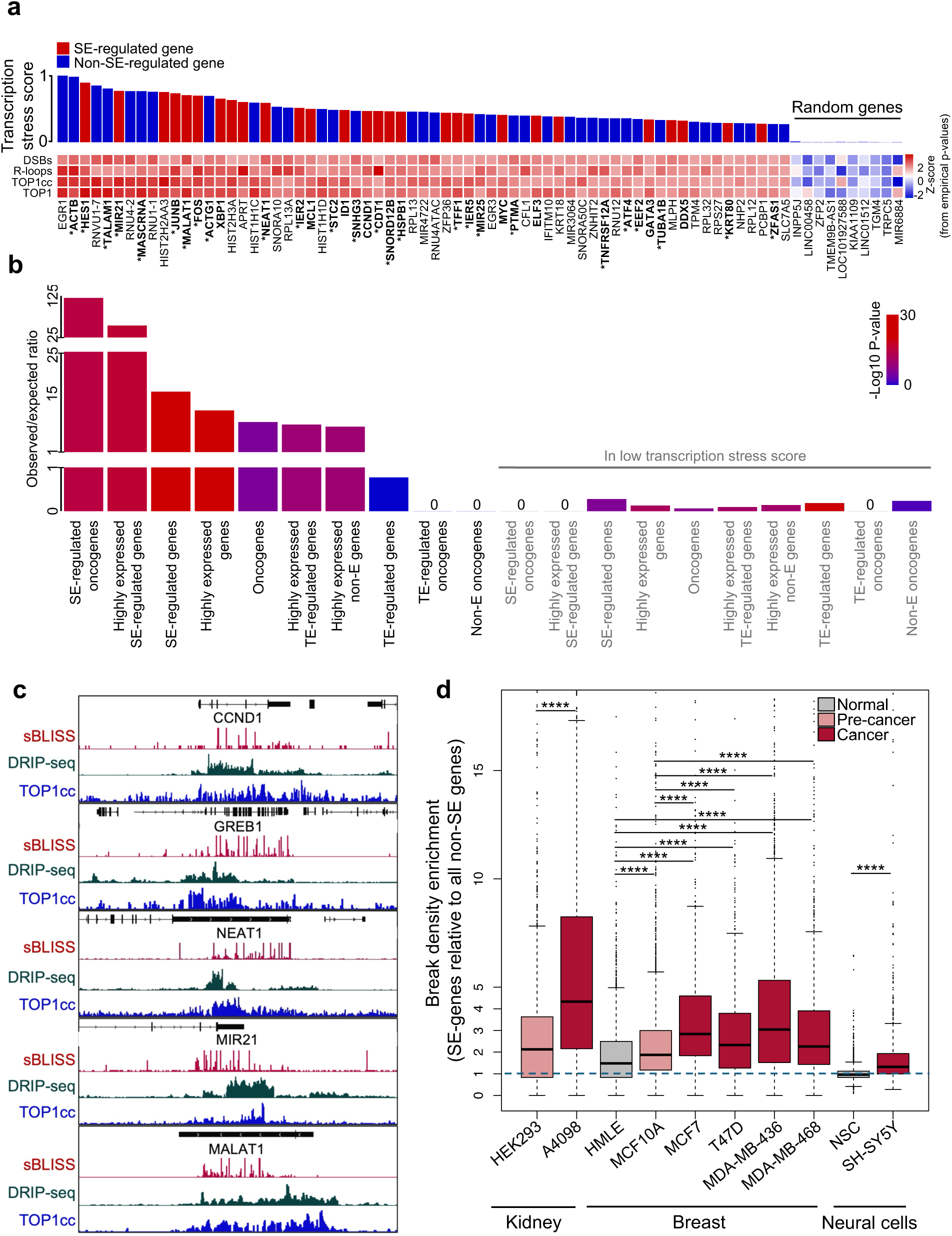
Super-enhancer-regulated oncogenes are major transcription stress sites. **a**, Transcription stress sites ranked by their scores based on four transcription stress parameters: TOP1, R-loops, TOP1cc, and DSBs in MCF7 cells (see methods). Genes with known oncogenic function in OncoKB are in bold and from other resources in bold with an asterisk (see table 1). **b,** The ratio of observed to expected numbers of genes in transcription stress sites. P-values were calculated using hypergeometric testing (upper-tailed for enrichment in the top score and lower-tailed for the bottom score). **c,** Genome browser snapshots depict DSBs, TOP1cc, and R-loops at selected transcription stress sites. **d,** A boxplot illustrates DSBs enrichment at the SE-regulated genes over other genes in each cell line (downloaded from SEdb^2.0^^31^). Enrichment was calculated by dividing the break density of each SE-regulated gene by the median break density of non-SE-regulated genes and the average values for replicates were plotted. In the boxplot, the center line represents the median; the box limits indicate the upper and lower quartiles; the whiskers denote 1.5X the interquartile range, while the dots represent outliers. P-values were computed using the Mann-Whitney U test, ****p < 0.0001.

**Table 1:** transcription stress sites genes and whether they have a known oncogenic function.

| Gene | Oncogenic function | Source |
| --- | --- | --- |
| EGR1 | no |  |
| ACTB | yes | Guo C. et al.,<sup>74</sup> |
| HES1 | yes | Wang J. et al.<sup>75</sup> |
| RNVU1-7 | No |  |
| TALAM1 | yes | Bermudez G. et al.<sup>76</sup> |
| MIR21 | yes | Zhu S. et al.<sup>77</sup> |
| RNU4-2 | no |  |
| MASCRNA | yes | Xie S. et al.<sup>78</sup> |
| RNU1-1 | no |  |
| HIST2H2AA3 | no |  |
| JUNB | yes | Ren F. et al.,<sup>79</sup> |
| MALAT1 | yes | Li Z. et al.<sup>80</sup> |
| FOS | yes | Ordway J. et al.<sup>81</sup> |
| ACTG1 | yes | Tang Y. et al.<sup>82</sup> |
| XBP1 | yes | OncoKB |
| HIST2H3A | No |  |
| APRT | No |  |
| HIST1H1C | No |  |
| NEAT1 | yes | Yu X. et al.<sup>83</sup> |
| SNORA10 | No |  |
| RPL13A | No |  |
| IER2 | yes | Kyjacova L. et al.<sup>84</sup> |
| MCL1 | yes | OncoKB |
| HIST1H1D | No |  |
| STC2 | yes | Qie S. et al.<sup>85</sup> |
| ID1 | yes | OncoKB |
| SNHG3 | yes | Xu b. et al.<sup>86</sup> |
| CCND1 | yes | OncoKB |
| CDT1 | yes | Wang S. et al.<sup>87</sup> |
| SNORD12B | yes | Zhao X. et al.<sup>88</sup> |
| HSPB1 | Yes | Liang Y. et al.<sup>89</sup> |
| RPL13 | No |  |
| MIR4722 | No |  |
| RNU4ATAC | No |  |
| ZFP36 | No |  |
| TFF1 | yes | Perry J. et al.<sup>90</sup> |
| IER5 | yes | Asano Y. et al.<sup>91</sup> |
| MIR25 | yes | El-Mezayen H. et al.<sup>92</sup> |
| EGR3 | No |  |
| MYC | yes | OncoKB |
| PTMA | yes | Zhu Y. et al.<sup>93</sup> |
| CFL1 | No |  |
| ELF3 | yes | OncoKB |
| IFITM10 | No |  |
| KRT18 | No |  |
| MIR3064 | no |  |
| SNORA50C | no |  |
| ZNHIT2 | No |  |
| TNFRSF12A | yes | Wang C. et al.<sup>94</sup> |
| RNU12 | No |  |
| ATF4 | yes | Wortel I. et al.<sup>95</sup> |
| EEF2 | yes | Oji Y. et al.<sup>96</sup> |
| GATA3 | yes | OncoKB |
| TUBA1B | yes | Wang Y. et al.<sup>97</sup> |
| MLPH | No |  |
| DDX5 | Yes | OncoKB |
| TPM4 | No |  |
| RPL32 | No |  |
| RPS27 | No |  |
| KRT80 | yes | Wei X. et al.<sup>98</sup> |
| NHP2 | No |  |
| RPL12 | No |  |
| PCBP1 | No |  |
| ZFAS1 | yes | He A. et al.<sup>99</sup> |
| SLC7A5 | no |  |

DSBs at these genes coincide with TOP1cc and R-loops (Fig. 1C). Analysis of sBLISS data for MCF7 cells depleted for TOP1 and R-loops shows that most of these genes have significantly fewer breaks upon TOP1 and R-loops depletion (Extended data Fig. 2A). Additionally, DSB are enriched at SE-regulated genes in both cycling and G1-synchronized cells (Extended data Fig. 2C-F), validating their transcriptional origin and independence from replication.

**Fig. 2:**
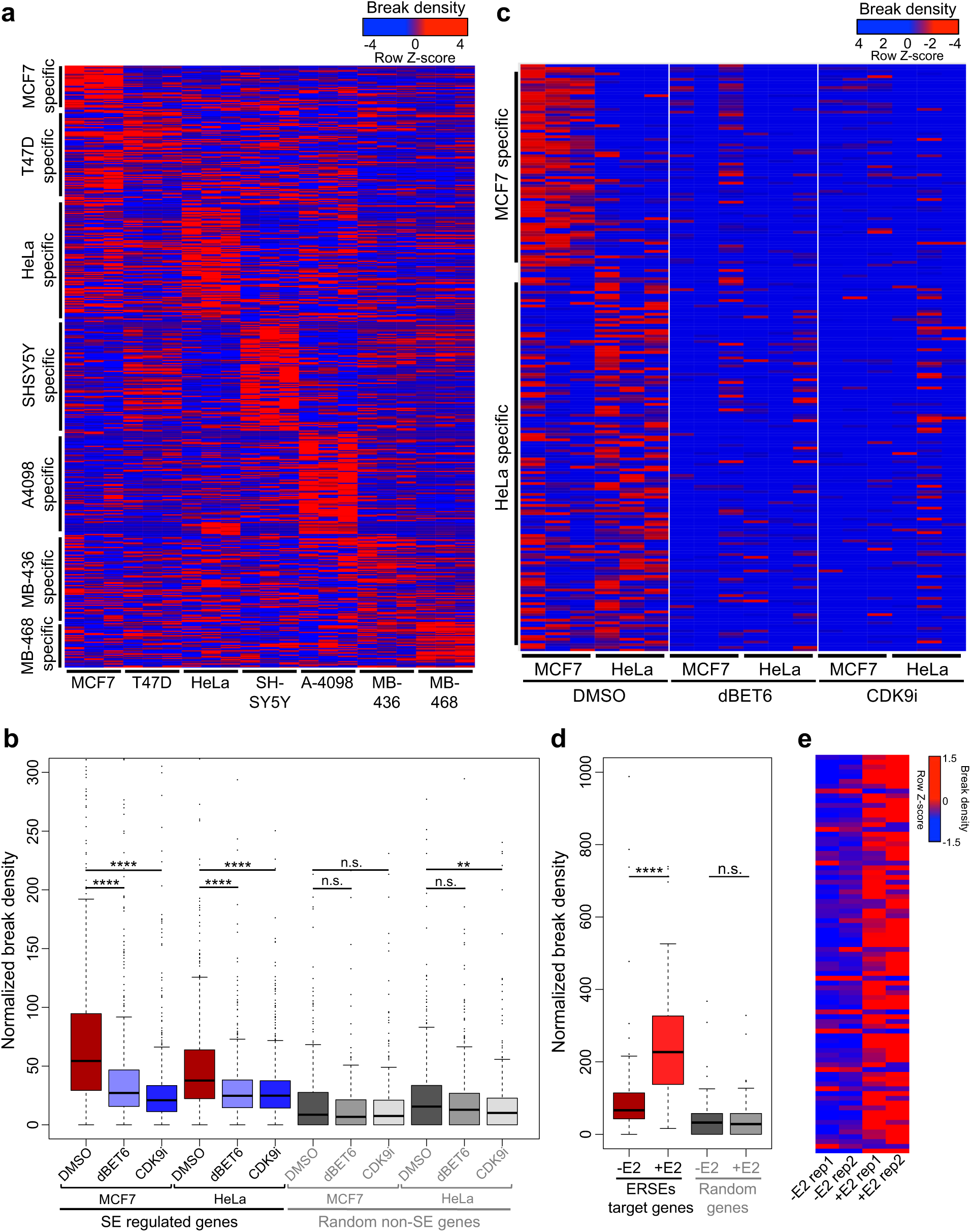
Super-enhancer landscape shapes transcription stress sites in cancer. **a**, Heatmap showing the normalized break density row Z-score for each cell line’s unique SE-regulated genes. For each cell line, SE-regulated genes were identified, common SE-regulated genes from other cell lines were excluded, and low to moderately expressed genes as well as genes with low break counts were filtered out. Each column corresponds to a biological replicate. Columns were normalized by median scaling. **b**, Boxplot illustrating the normalized break density of cells treated with DMSO (control), dBET6, or CDK9i (100nM for 4 hours each) for SE-regulated genes in each cell line or for randomly selected genes. values are averaged for three biological replicates. **c**, Heatmap depicting the normalized break density row Z-score for each cell line’s unique SE-regulated genes following treatment with DMSO (control), dBET6, or CDK9i. Each column corresponds to a biological replicate. Each sample was normalized by downsizing to 270000 random breaks. **d**, Boxplot demonstrating the normalized break density of cells treated with vehicle (control) or estradiol (E2, 100nM for 1 hour) for significantly affected (p < 0.05) estrogen receptor-occupied SE (ERSE) target genes or for randomly selected genes. **e**, Heatmap representing the normalized break density row Z-score of significantly affected ERSE target genes. Boxplot features include: center line representing median; box limits denoting upper and lower quartiles; whiskers extending to 1.5x the interquartile range; and dots indicating outliers. P-values were computed using the Mann-Whitney U test, with n.s. indicating p > 0.05, ** indicating p < 0.01, and **** indicating p < 0.0001.

The relationship between transcription and DSBs was previously established^13–15^. However, our results suggest that this association may be more assertive in SE-regulated genes. This prompted us to further investigate the discrepancies in the relationship between transcription and DSBs in SE-regulated versus non-SE-regulated genes. Interestingly, the strong correlation between gene expression and DSBs is only observed in SE-regulated genes and not in others (Extended Data Fig. 2G-I). Moreover, highly expressed genes enriched with DSBs showed significantly more enrichment in TOP1, TOP1cc, and R-loops than similarly expressed genes lacking DSBs (Extended Data Fig. 3A-E). Additionally, SE-regulated genes were significantly more prevalent in highly expressed genes enriched with DSBs (Extended Data Fig. 3F). These findings indicate that the association between transcription and DSBs is primarily attributed to SE-regulated genes.

Since SE activity is significantly more potent in cancer cells than in normal cells, we next aimed to test whether the enrichment of DSBs at SE-regulated genes is a phenomenon specific to cancer. To investigate this, we mapped the Breakome in 10 different cell lines, which included three normal cell lines, one pre-malignant cell line, and six cancer cell lines. These results indicate that SE-regulated genes are more enriched with DSBs in cancer cells than in their normal counterparts (Fig. 1D), demonstrating the specificity of transcriptional stress to active SE-regulated oncogenes in cancer cells.

### The super-enhancer landscape shapes transcription stress sites in cancer

The landscape of SEs is cell-type specific, enhancing the transcription of different genes based on cell identity^12^. If SEs cause transcription stress in their target genes, one would predict that various cancer cell types exhibit distinct transcription stress sites reflective of their SE landscapes. To test this, we mapped the Breakome of multiple cancer cell lines. For each cell line, SE-regulated genes were identified, common SE-regulated genes from other cell lines were excluded, and low to moderately expressed genes, as well as genes with low break counts, were filtered out. Interestingly, SE-regulated genes were enriched in DSBs, predominantly in the cell type where they are regulated by an SE (Fig. 2A). This pattern was also apparent when comparing cell types with very similar characteristics (MCF7 and T47D, both of which are ER+ luminal A breast cancer cell types), yet exhibit differing SE landscapes, suggesting that transcription stress sites correspond to the SE landscape in cancer.

If SE activity indeed causes the accumulation of DSBs at transcription stress sites, then inhibiting their activity would abolish SE-mediated DSB accumulation, while conversely, activating it would enhance DSB accumulation at target genes. To test this hypothesis, we employed two strategies. First, we inhibited SE activity in MCF7 and HeLa cells through two approaches, using the BRD4 degron dBET6 and the CDK9 inhibitor (CDK9i) NVP-2, both of which have been shown to lead to SE collapse and repress SE-mediated transcription^32,33^. Examining the breakome after a 4-hour incubation with these inhibitors, which effectively suppressed SE-regulated transcriptional activity as measured by spike-in real-time quantitative PCR (RT-qPCR) (Extended data Fig. 3G-H), revealed that SE inhibition significantly reduced DSBs at SE-regulated genes in both cell lines (Fig. 2B). Furthermore, SE inhibition using either dBET6 or CDK9i eliminated the cell-type-specific effect of the SE landscape on the breakome (Fig. 2C).

**Figure 3.**
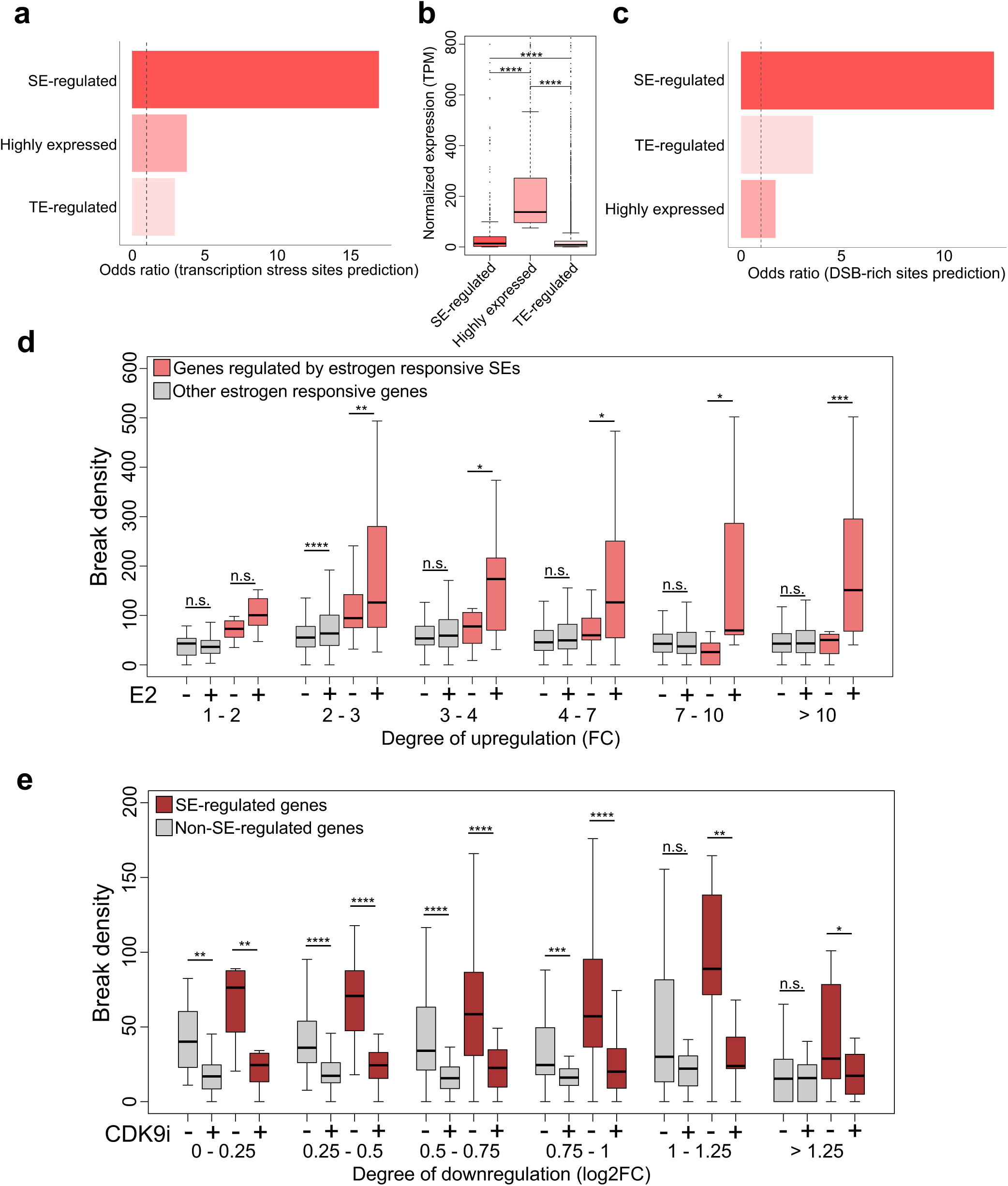
Specificity of transcription stress at super-enhancer (SE)–regulated genes. **a**, Multivariate LASSO regression identifying predictors of transcription stress site locations in 100kb genomic bins. Odds ratios are shown. **b,** Boxplot showing normalized expression (TPM) of SE-regulated, highly expressed, and TE-regulated genes. Despite lower expression. **c,** LASSO regression for prediction of DSB-rich genomic sites. the genome was binned in 10kb bins, and the top 5% bind in break density were assigned as DSB-rich. **d,** Boxplot showing the break density in estrogen-responsive genes upon E2 treatment, grouped by degree of transcriptional upregulation (fold change, FC). Non-SE genes were randomly downsampled to match SE gene numbers for comparable P-values. **e,** Boxplot showing the break density in estrogen-responsive genes following CDK9 inhibition (CDK9i), grouped by degree of transcriptional downregulation (log2 fold change). Non-SE genes were randomly downsampled to match SE gene numbers for comparable P-values. Boxplot features include: center line representing the median; box limits denoting upper and lower quartiles; whiskers extending to 1.5x the interquartile range; and dots indicating outliers. P-values in boxplots were calculated using the Mann-Whitney U test in b, and Wilcoxon paired test in D and E. n.s. p >0.05, *p < 0.05, **p < 0.01, ***p < 0.001, ****p < 0.0001.

Secondly, we utilized the fact that MCF7 cells are estrogen receptor-positive and that many of their SEs are estrogen-responsive and analyzed the breakome of these cells before and after estrogen addition. We specifically focused on genes regulated by estrogen receptor-occupied SE-regulated genes (ERSE-genes), which were previously characterized^34^. Consistently, ERSE induction significantly increased DSBs in target genes but not random genes (Fig. 2 D-E), indicating that SE activity promotes the accumulation of DSBs at SE-regulated genes. Overall, these data demonstrate that the landscape of SE activity in cancer cells shapes the landscape of transcription stress in a cell-type and cancer-type-specific manner.

### Specificity of transcription stress at super-enhancer-regulated genes

While multiple studies have demonstrated the formation of DSBs as a consequence of transcription in general^35–37^, our results indicate that this phenomenon is markedly amplified when transcription is driven by SEs (Fig. 1B; Extended Data Fig. 1E–F). To further support this observation, we employed two complementary approaches.

First, we examined predictors for the genomic location of transcription stress sites. Using SE-regulated genes, typical enhancer (TE)-regulated genes, and general highly expressed genes as predictors, multivariate LASSO regression revealed that SE regulation was the strongest predictor of transcription stress sites (odds ratio = 12.61; Fig. 3A), despite the highly expressed gene group exhibiting the highest transcriptional levels overall (Fig. 3B). Notably, SE regulation was also the best predictor for genomic regions enriched in DSBs (Fig. 3C).

Second, to experimentally assess whether transcriptional activation through SEs promotes greater transcription stress and DSB accumulation, we performed a deeper analysis on MCF7 breakome before and after estradiol treatment. Estrogen activates transcription via both promoter-proximal binding and estrogen-responsive SEs^38^, enabling us to compare the impact of these two mechanisms. We grouped genes by their level of transcriptional upregulation (based on GRO-seq fold change) and further subdivided each group into SE-regulated and non-SE-regulated categories. Consistently, SE-regulated genes accumulated significantly more DSBs than their non-SE counterparts. This effect was especially prominent in the highest induction group (fold change > 10; Fig. 3D). A similar pattern was observed upon CDK9 inhibition, where SE-regulated genes exhibited a significantly greater reduction in DSBs compared to non-SE-regulated genes with matched transcriptional downregulation (measured by RNA-seq) (Fig. 3E).

Together, these results demonstrate that SE-mediated transcription is a major determinant of transcription stress site formation and leads to amplified DSB accumulation compared to non-SE transcriptional activity, even at equivalent expression levels.^38^

### γH2AX marks transcription stress sites

Since DSBs are transcription-blocking lesions^28^, transcription stress sites must be rapidly and efficiently repaired to maintain high expression levels. γH2AX is a marker for DDR activation and is among the first responders to DSBs, as it is phosphorylated within a few minutes after DSB formation^39^. To further explore the role of DDR activation in the signaling of DSB repair at transcription stress sites, we conducted ChIP-seq for γH2AX on untreated MCF7 cells and analyzed high-quality available γH2AX ChIP-seq data for HeLa and IMR90 cell lines.

In both MCF7 and HeLa cells, γH2AX signal is predominantly localized to gene bodies, whereas in IMR90 cells the distribution is more balanced between genic and intergenic regions (Extended Data Fig. 4A). This pattern is consistent with a greater burden of transcription-associated stress in cancer cells compared to non-transformed cells. Moreover, γH2AX is distributed across the gene bodies of top targets (Extended Data Fig. 4B-D). Ranking genes based on their γH2AX enrichment revealed that most exhibit relatively low γH2AX levels, while a subset displays high to very high γH2AX density (Fig. 4A). This finding indicates a non-uniform distribution of γH2AX, with preferential enrichment at specific genes. Interestingly, all transcription stress sites are among the top 8% of γH2AX targets (Fig. 4A-B). This enrichment in the top γH2AX genes was much less for the general high DSB genes (Fig. 4C). Additionally, top γH2AX targets are enriched in SE-regulated genes across each respective cell line (P= 2.3e^-41^, 2.9e^-39^, 8.5e^-8^, for MCF7, HeLa, and IMR90 respectively; hypergeometric test) (Fig. 4D-F).

**Fig. 4:**
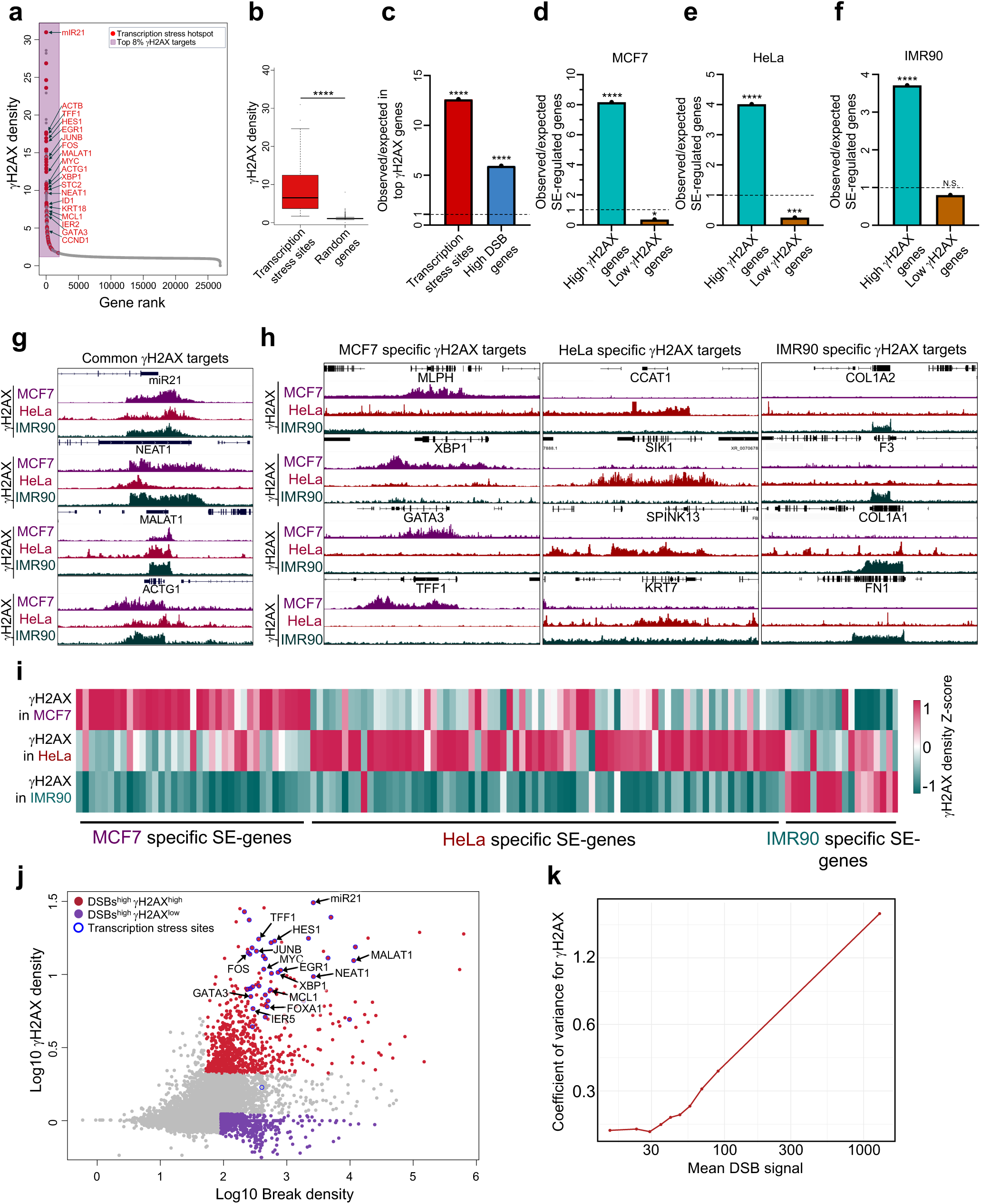
gH2AX marks transcription stress sites. **a**, Ranked plot of genes in MCF7 cells based on γH2AX density. γH2AX density per gene was computed using bigWigAverageOverBed, which calculates the average signal from the γH2AX bigWig track over annotated gene intervals. **b**, Boxplot showing γH2AX density for transcription stress sites and randomly selected genes. **c**, The ratio of the observed to the expected number of transcription stress sites or top 1000 break-prone genes in MCF7 among the top γH2AX enriched genes. **d**-**f**, The ratio of observed to expected numbers of SE-regulated genes in the top and bottom 300 genes enriched in γH2AX for each cell line. P-values were determined using the hypergeometric test: n.s. p > 0.05, *p < 0.05, ***p < 0.001, ****p < 0.0001. **g,** Genome browser snapshots of representative common γH2AX targets across MCF7, HeLa, and IMR90 cell lines. **h**, Genome browser snapshots of representative unique γH2AX targets in MCF7, HeLa, and IMR90 cell lines. **i**, Heatmap displaying the γH2AX density column Z-score for each cell line’s unique SE-regulated genes. For each cell line, SE-regulated genes were identified, common SE-regulated genes with other cell lines, and genes with low γH2AX counts were filtered out. **j**, Scatter plot showing the correlation between break density and γH2AX density for each gene. **k,** Coefficient of variation (CV) of γH2AX density as a function of mean DNA double-strand break (DSB) signal. Genes were binned into deciles based on their DSB signal intensity (log-transformed), and for each bin, the mean γH2AX density, standard deviation, and CV (standard deviation divided by mean) were calculated. The x-axis shows the mean DSB signal per bin on a log scale, and the y-axis shows the CV of γH2AX density within each bin. Boxplot features include: center line representing the median; box limits denoting upper and lower quartiles; whiskers extending to 1.5x the interquartile range; and dots indicating outliers. P-values in boxplots were calculated using the Mann-Whitney U test, ****p < 0.0001.

The observed γH2AX enrichment across the three cell lines was consistent for several oncogenes, including miR21, NEAT1, MALAT1, and ACTG1 (Fig. 4G). Interestingly, many prominent γH2AX targets were unique to specific cell lines, most with established activity in their respective cell types (Fig. 4H). For example, two top γH2AX targets in MCF7, TFF1 and GATA3, are well-known oncogenic transcription factors primarily active in breast cancer and were exclusively enriched in γH2AX in MCF7 cells. KRT7 and CCAT1, which are overexpressed and linked to poor prognosis in cervical cancer, are enriched in γH2AX only in HeLa cells. Consequently, we questioned whether these variations in γH2AX enrichment stem from cell-type-specific SEs. To investigate this, we compared γH2AX enrichment of top targets that are regulated by a cell-type specific SE in each respective cell line. Indeed, SE-regulated γH2AX targets are predominantly enriched in the cell line where they are regulated by an SE (Fig. 4 I), underscoring that γH2AX marks cell type-specific transcriptional stress sites.

### γH2AX levels vary significantly across genes enriched with high endogenous DSBs

Next, we examined the correlation between DSBs and γH2AX in genes. Interestingly, analyzing the correlation between each gene’s breaks and its γH2AX density revealed high variability of γH2AX among genes with high DSBs (Fig. 4 J-K and Extended Data Fig. 5A-E). Some genes enriched with breaks are marked by γH2AX while others are not, in both MCF7 and HeLa cell lines (Extended Data Fig. 5F-G). This discrepancy in γH2AX enrichment at various DSB-enriched genes might suggest that regions enriched with endogenous DSBs in the genome are signaled with varying efficiency and potentially repaired unevenly, depending on their context.

### γH2AX preferentially marks endogenous DSBs of active chromatin

Although genes exhibit a moderate correlation between DSB density and γH2AX density (Spearman’s correlation = 0.57), non-linear regression analysis using Generalized Additive Models (GAM) indicates that the transcription stress score explains a greater proportion of the variance in γH2AX levels (R² = 0.73) than any individual component (Fig. 5A and Extended data Fig. 6). To further investigate the underlying factors contributing to variability in γH2AX at high DSBs genes, we defined two groups of genes with similar break density but different γH2AX levels (Fig. 4J): DSBs^high^ γH2AX^high^ genes and DSBs^high^ γH2AX^low^ genes. DSBs^high^ γH2AX^high^ genes are more expressed and enriched with transcription stress markers (TOP1, TOP1cc, and R-loops) (Extended data Fig. 7A-E). Furthermore, 64 of the 65 transcriptional stress sites were among DSBs^high^ γH2AX^high^ genes (Fig. 4J and Fig. 5B). This group also included all SE-regulated transcription stress sites and is significantly enriched with active SE-regulated genes compared to DSBs^high^ γH2AX^low^ genes, which lack active SE-regulated genes and transcriptional stress hotspots (Fig. 5C and Extended data Fig. 7F).

**Fig. 5:**
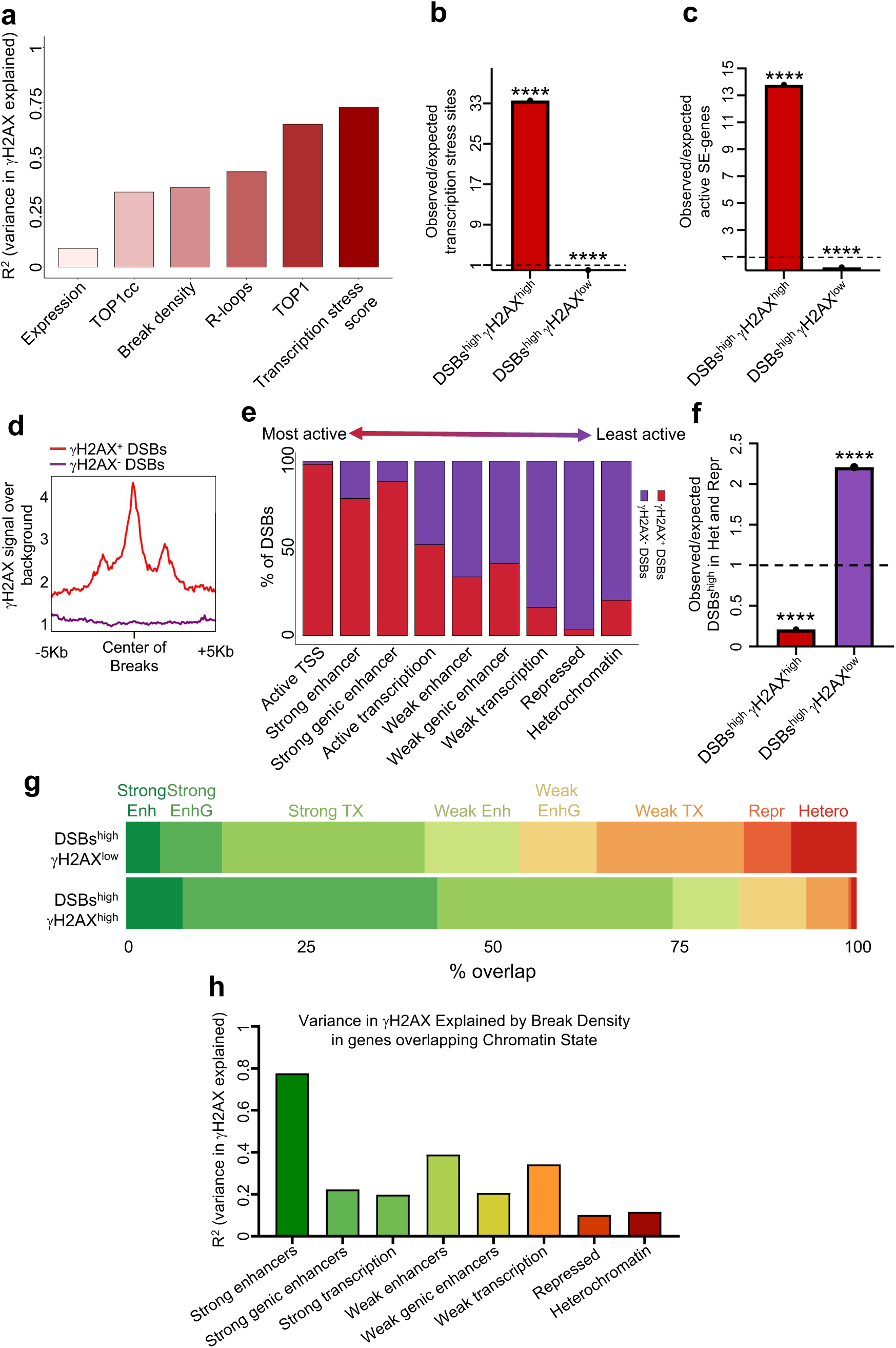
γH2AX preferentially marks endogenous DSBs of active chromatin. **a**, Barplot showing R² values for the different variables tested regarding their effect on γH2AX variance across all genes. **b**-**c**. The ratio of observed to expected number of transcription stress sites (**b**) and active SE-regulated genes (**c**). **d**, γH2AX signal at the two defined sets of DSBs. **e**. The percentage of DSBs enriched with γH2AX (γH2AX^high^ DSBs) or lacking γH2AX (γH2AX^low^ DSBs) in each chromatin state. Chromatin states were ordered based on what is typically known about their transcriptional activity^73^. **f**, The ratio of observed to expected numbers of high DSB genes overlapping heterochromatin and repressed regions in both gene groups in MCF7 cells. **g**. The percentage of overlaps between each gene group and chromatin states in MCF7. For each group, the number of overlaps was counted for each chromatin state and then divided by the total number of overlaps. **h**. A barplot showing R² values for the effect of DSBs on γH2AX variance for genes overlapping different chromatin states. P-values were determined using the hypergeometric test: ****p < 0.0001.

**Fig. 6:**
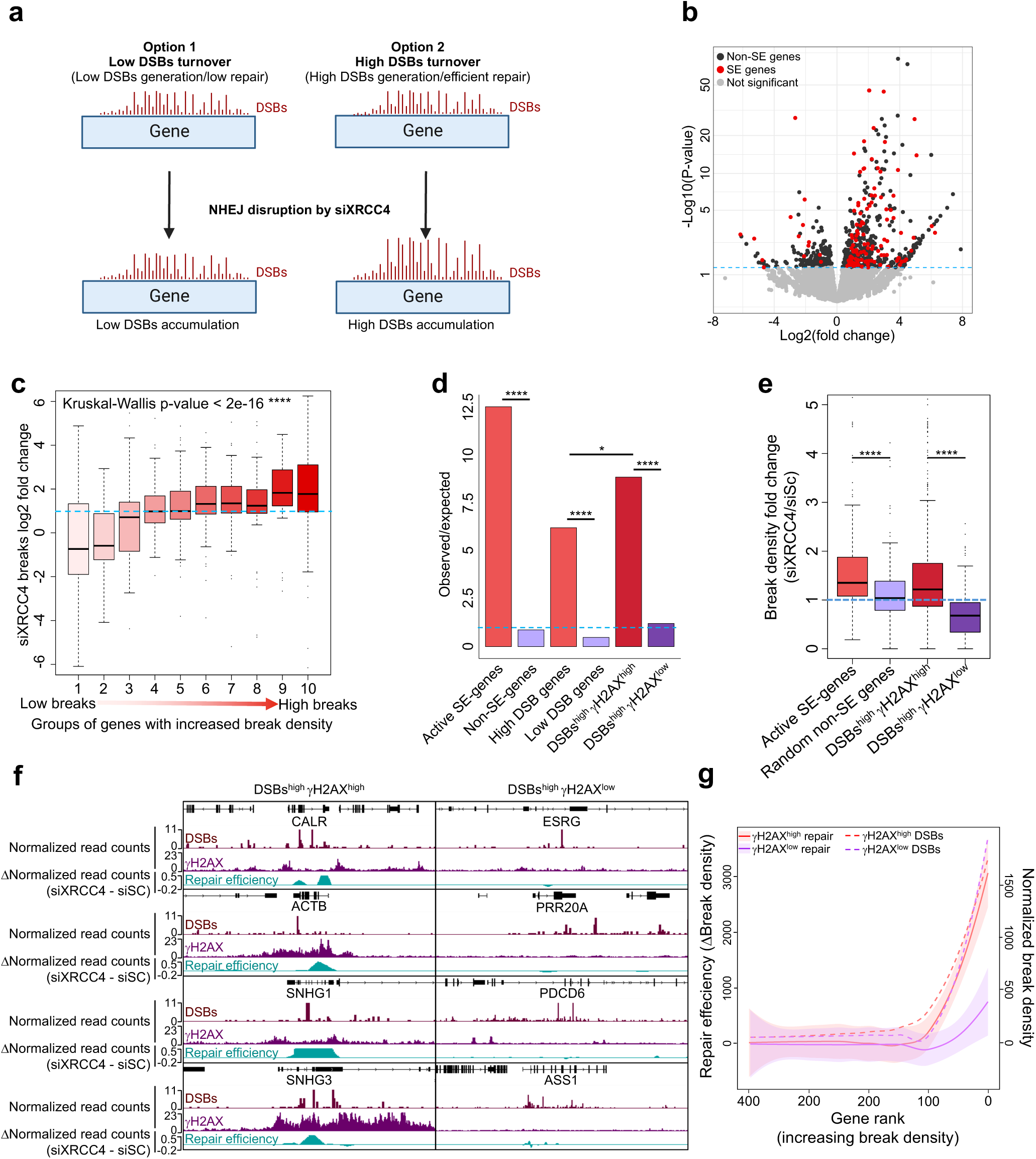
Genes marked with γH2AX show high DSB turnover and repair efficiency. **a**. Schematic representation of the rationale behind the experiment to estimate the rate of endogenous DSB repair and turnover. **b**. Volcano plot depicting the results of the differential expression analysis of DSBs following XRCC4 KD, assessed using the DESeq2 algorithm. The experiment was performed on two biological replicates per condition. **c**. Boxplot illustrating the log2FC of break density upon XRCC4 KD for significantly affected genes stratified according to their baseline break density into ten groups. **d**. The ratio of observed to expected number of genes in our defined gene sets. P-values were calculated using Fischer exact test. **e**. Boxplot of break density fold change. P-values were calculated using the Mann-Whitney U test. **f**. Genome browser snapshots of DSBs^high^ γH2AX^high^ and DSBs^high^ γH2AX^low^ gene examples, showing DSBs, γH2AX, and repair efficiency. Repair efficiency was estimated by pooling replicates, tiling the genome into 100bp tiles, subtracting the break score of each tile in the siXRCC4 sample from siSc (control), and smoothing the segments through sliding window averaging. **g**. Line plot for repair efficiency (log2FC after XRCC4 KD) for DSBs^high^ γH2AX^high^ versus DSBs^high^ γH2AX^low^ genes ordered by their break density. Lines were generated using LOESS smoothing. Boxplot: center line indicates median; box limits represent upper and lower quartiles; whiskers extend to 1.5x interquartile range; dots indicate outliers. *p < 0.05, ****p < 0.0001.

**Fig. 7:**
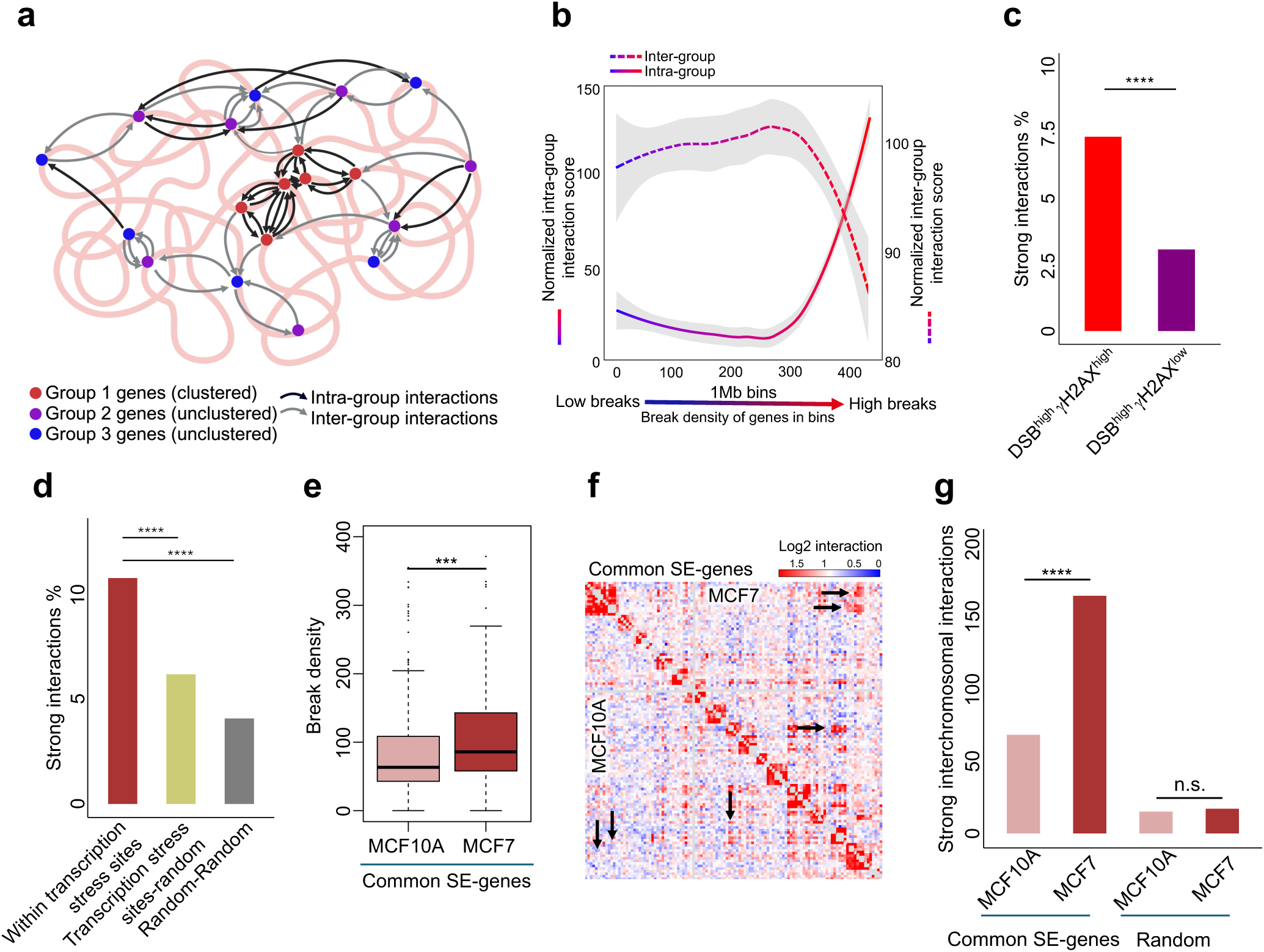
Efficiently repaired transcriptional DSBs cluster in the 3D genome. **a**. Schematic illustrating inter– and intra-group interactions and their use to determine the clustering of genes with a common characteristic (high DSB density in this instance). **b**. Line plot depicting inter-group (dashed line) and intra-group (solid line) genomic bins arranged according to the break density of their overlapping genes. Lines were created using LOESS smoothing, with shaded areas representing the 95% confidence intervals. **c**. Percentage of strong interactions within genomic bins containing DSBs^high^ γH2AX^high^ compared to bins with DSBs^high^ γH2AX^low^ genes. Strong interactions are defined as a log2 interaction score > 2. The percentage is calculated by dividing the number of strong interactions by the total number of interactions. P-values were calculated by Chi-squared test. **d**. Percentage of strong interactions calculated as in (**c**). **e**. Boxplot reflecting break density of common SE-regulated genes between MCF7 and MCF10A cell lines. P-values were calculated by Mann-Whitney U test. **f**. Heatmap showing Log2 interaction scores for genomic bins containing common SE-regulated genes from both MCF7 and MCF10A cell lines. Arrows indicate distinct sites of variation in interactions. **g**. Count of interchromosomal strong interactions shown in the heatmap from (**f**). Random bins serve as a reference. Boxplot: center line indicates median; box limits represent upper and lower quartiles; whiskers extend to 1.5x interquartile range; dots indicate outliers. P-values were calculated using chi-square test in (**c**-**d**) and (**g**), and Wilcoxon paired test in (e). ***p < 0.001, ****p < 0.0001.

Chromatin context was previously shown to impact the phosphorylation of H2AX, specifically, heterochromatin was shown to be refractory to H2AX phosphorylation^40^. Therefore, we wondered whether the observed DSBs^high^ γH2AX^low^ genes are close to, or within, heterochromatic regions. To gain further insights into whether γH2AX preferentially marks endogenous DSBs based on the chromatin context, we defined two groups of individual DSB sites across the genome according to their γH2AX signal (Fig. 5D and Extended data Fig. 7G) and mapped them to the different chromatin states of MCF7 and HeLa. γH2AX^high^ DSBs were biased toward active chromatin states, while γH2AX^low^ DSBs were biased toward silent regions (Fig. 5E and Extended data Fig. 7H). These results made us question whether the differential γH2AX marking at DSB-enriched genes was due to their chromatin context. Indeed, the DSBs^high^ γH2AX^low^ group, but not the DSBs^high^ γH2AX^high^ group, is enriched with DSB-enriched genes that overlap with heterochromatin and repressed chromatin states (Fig. 5F and Extended data Fig. 7I). Additionally, an analysis of the chromatin state contexts of DSBs^high^ γH2AX^high^ and DSBs^high^ γH2AX^low^ in both MCF7 and HeLa showed that DSBs^high^ γH2AX^high^ genes primarily overlap with chromatin regions associated with strong transcriptional activity. Conversely, DSBs^high^ γH2AX^low^ has a higher overlap with heterochromatin, repressed, and weak transcription chromatin states (Fig. 5G and Extended data Fig. 7J).

Our results illustrate that DSBs alone do not fully account for the variation in γH2AX across genes (Fig. 5A). Conducting GAM on genes categorized by their chromatin context revealed that DSBs primarily account for variation in γH2AX in genes that overlap with strong enhancer regions and least in genes that overlap with repressed or heterochromatin regions (Fig. 5H). This suggests that breaks and γH2AX are most closely associated with genes spanning strong enhancer elements.

Collectively, these results indicate that γH2AX preferentially marks endogenous DSBs in active chromatin compared to inactive chromatin. This differential marking may influence variations in the repair efficiency of distinct DSBs based on their chromatin context and γH2AX signalling.

### Genes marked with γH2AX exhibit high DSB turnover and repair efficiency

sBLISS, like other methods used to map DSBs, captures a snapshot of the DSB landscape at the moment of fixation^41–45^. Therefore, a gene that shows high levels of DSBs could experience frequent DSB generation and repair cycles (high DSB turnover) or infrequent DSB generation and repair cycles (low DSB turnover). To estimate the rate of endogenous DSB repair across the genome, we mapped DSBs in HeLa cells after disrupting the two major DSB repair pathways, non-homologous end-joining (NHEJ)^45^ and homologous recombination (HR), by using siRNAs that target XRCC4 and RAD51, respectively (Extended Data Fig. 8A). We applied DESeq2 ^46^ to the DSB count matrices to identify genes with significant DSB accumulation following knockdown of each factor. Hierarchical clustering of these genes revealed three distinct response groups: Cluster 1 genes were primarily affected by XRCC4 depletion, Cluster 2 by both XRCC4 and RAD51 (with a stronger effect from RAD51), and Cluster 3 mainly by RAD51 depletion (Extended Data Fig. 8B). Importantly, no cluster was exclusively dependent on RAD51, consistent with previous reports that XRCC4 localizes to both RAD51-bound and unbound breaks^47^.

Further analysis revealed that all three clusters are associated with transcription stress, with Cluster 1 (XRCC4-dependent genes) showing the strongest enrichment (Extended Data Fig. 8C). These genes also exhibit the highest baseline break density (Extended Data Fig. 8D), suggesting that NHEJ is particularly engaged at the most DSB-prone, transcriptionally stressed loci. Collectively, these findings support the involvement of both NHEJ and HR in repairing transcription-associated DSBs, while indicating a prominent role for XRCC4-mediated repair at the most vulnerable genomic sites. Based on this, we proceeded to quantify genome-wide DSB turnover using XRCC4 knockdown–induced break accumulation as a proxy for repair efficiency.

Regions that are frequently repaired should accumulate excessive DSBs due to repair impairment. In contrast, regions with inherently low repair rates should show minimal DSB accumulation (Fig. 6A). 429 genes showed significant DSB accumulation upon XRCC4 KD (Fig. 6B). Stratifying these genes by their baseline DSB density revealed a trend in which groups with higher baseline DSB density showed progressively greater impact following XRCC4 KD (Fig. 6C), supporting the notion that DSB-prone genes are repaired more effectively. Enrichment analysis indicated a strong enrichment of DSBs^high^ γH2AX^high^ genes (Fig. 6D), along with the top DSB-enriched and active SE-regulated genes. In contrast, DSBs^high^ γH2AX^low^ genes and the bottom DSB-enriched and non-SE-regulated genes were not significantly enriched. Directly examining the effects of repair impairment demonstrated that NHEJ impairment leads to the most pronounced accumulation of DSBs at DSBs^high^ γH2AX^high^ and SE-regulated genes, compared to DSBs^high^ γH2AX^low^ and non-SE genes (Fig. 6EE). To investigate further how γH2AX patterns influence the repair turnover of endogenous DSBs, we used the accumulation of DSBs following repair impairment as a proxy for repair efficiency (Fig. 6F). Within DSBs^high^ γH2AX^high^ genes, repair efficiency markedly increases with higher DSB levels; however, within DSBs^high^ γH2AX^low^ genes, this relationship of repair efficiency is significantly weaker (Fig. 6G). These findings suggest two distinct types of endogenous DSBs: γH2AX-marked DSBs, which exhibit high turnover, and poorly γH2AX-marked DSBs, which exhibit low turnover.

### Efficient repair at transcriptional stress sites is crucial for SE-regulated oncogene expression

High γH2AX marking and repair efficiency at transcription stress sites under physiological conditions suggest that these regions rely on DSB repair to sustain their elevated transcriptional levels. To test this, we inhibited DNA-PKcs, a core component of the canonical NHEJ pathway, using a selective small-molecule inhibitor (DNA-PKci) in MCF7 cells. We then performed sBLISS to map genome-wide DSBs and spike-in RT-qPCR to quantify transcript levels. To assess genome-wide DSB accumulation, we divided the genome into 10 kb bins, quantified DSBs per bin, and used DESeq2 to identify bins with significant DSB enrichment in DNA-PKci-treated cells. We then overlapped significantly differential bins with annotated gene regions to determine whether specific gene classes were disproportionately affected. Indeed, significantly upregulated bins were strongly enriched at SE-regulated genes and not at non-SE-regulated genes (Extended Data Fig. 9A–b).

Measuring mRNA levels of the top γH2AX marked SE-regulated genes upon DNA-PKci indicated a significant decrease in their expression levels (Extended data Fig. 9C). To determine whether the observed transcriptional reduction was due to repair failure rather than secondary signaling effects, we also knocked down XRCC4 under the same conditions as in Fig. 6. This similarly led to downregulation of highly γH2AX-marked SE-regulated genes that accumulated DSBs (Extended Data Fig. 9D). Together, these results demonstrate that efficient DSB repair via the NHEJ pathway is required to sustain the transcription of SE-regulated oncogenes.

### Efficiently repaired transcriptional DSBs cluster in the 3D genome

Induced DSBs cluster in repair foci^48,49^, especially when introduced within active genes^49^. We hypothesized that endogenous transcription-associated DSBs might also cluster within the 3D genome as a mechanism to enhance repair. To investigate this, we analyzed high-quality Hi-C data generated from MCF7 cells^50^.

To quantify the clustering of genes associated with break density, we introduced two Hi-C metrics that measure how a group of genes with a common characteristic (e.g., break density) is clustered (Fig. 7A): intra-group interactions, which represent the sum of genomic interaction scores between genes within the same group, and inter-group interactions, the sum of genomic interaction scores between genes from different groups. A clustered group of genes is expected to exhibit higher intra-group interactions and lower inter-group interactions compared to a non-clustered group (Fig. 7A). To investigate whether DSB-enriched genes exhibit clustering, genomic bins were arranged according to the break densities of their constituent genes and categorized into nine groups based on break density. Indeed, bins containing DSB-enriched genes demonstrate higher intra-group interactions and lower inter-group interactions than bins with low DSB genes (Fig. 7B), indicating the clustering of high DSB genes.

The percentage of strong interactions among DSBs^high^ γH2AX^high^ genes is significantly higher than that among DSBs^high^ γH2AX^low^ genes (Fig. 7C and Extended data Fig. 9A). Additionally, transcription stress sites exhibit a substantially higher percentage of strong interactions compared to random sites with similar chromosomal distribution (Fig. 7 D and Extended data Fig. 9B), suggesting that efficiently repaired transcription stress sites are clustered together. To avoid the confounding effects of SE regulation on 3D interactions, we analysed Hi-C data from the MCF10A cell line, which was conducted parallel to MCF7^50^. MCF10A is a pre-cancerous cell line that shares many common SE-regulated genes with MCF7 cells. However, these common genes are significantly more enriched in DSBs within MCF7 cells (Fig. 7E). This enabled a comparison of the same regions with the same SE-regulation status but differing DSB enrichment. Indeed, bins containing common SE-regulated genes, as opposed to random regions, showed significantly higher strong interchromosomal interactions in MCF7 cells than in MCF10A cells (Fig. 7F-G), indicating that transcription stress sites undergo increased clustering.

### The repair of transcription stress sites is mutagenic

Since transcription stress sites are more likely to be in close proximity (Fig. 7), we suspected that recurrent DSBs at these sites could serve as mutational hotspots. To explore this, we downloaded and analysed mutational data from the COSMIC database^51^, which contains extensive mutational data on human cancers. While mutations within coding regions are subject to selective pressures based on their impact on protein function and cellular fitness ^52,53^, non-coding mutations—particularly those in intronic regions—are less constrained by selection and can provide a more direct readout of a region’s intrinsic susceptibility to mutation. To minimize the confounding effects of selection, we focused on non-coding intronic mutations in breast cancer and intersected them with our breakome data from the breast cancer cell line MCF7. Grouping genes by their DSB or γH2AX density revealed that genes with a higher break or γH2AX density exhibit greater intronic mutation density (Fig. 8A-C). This association suggests that the repair of transcription-associated DSBs is inherently error-prone, leading to the accumulation of mutations over time.

**Fig. 8:**
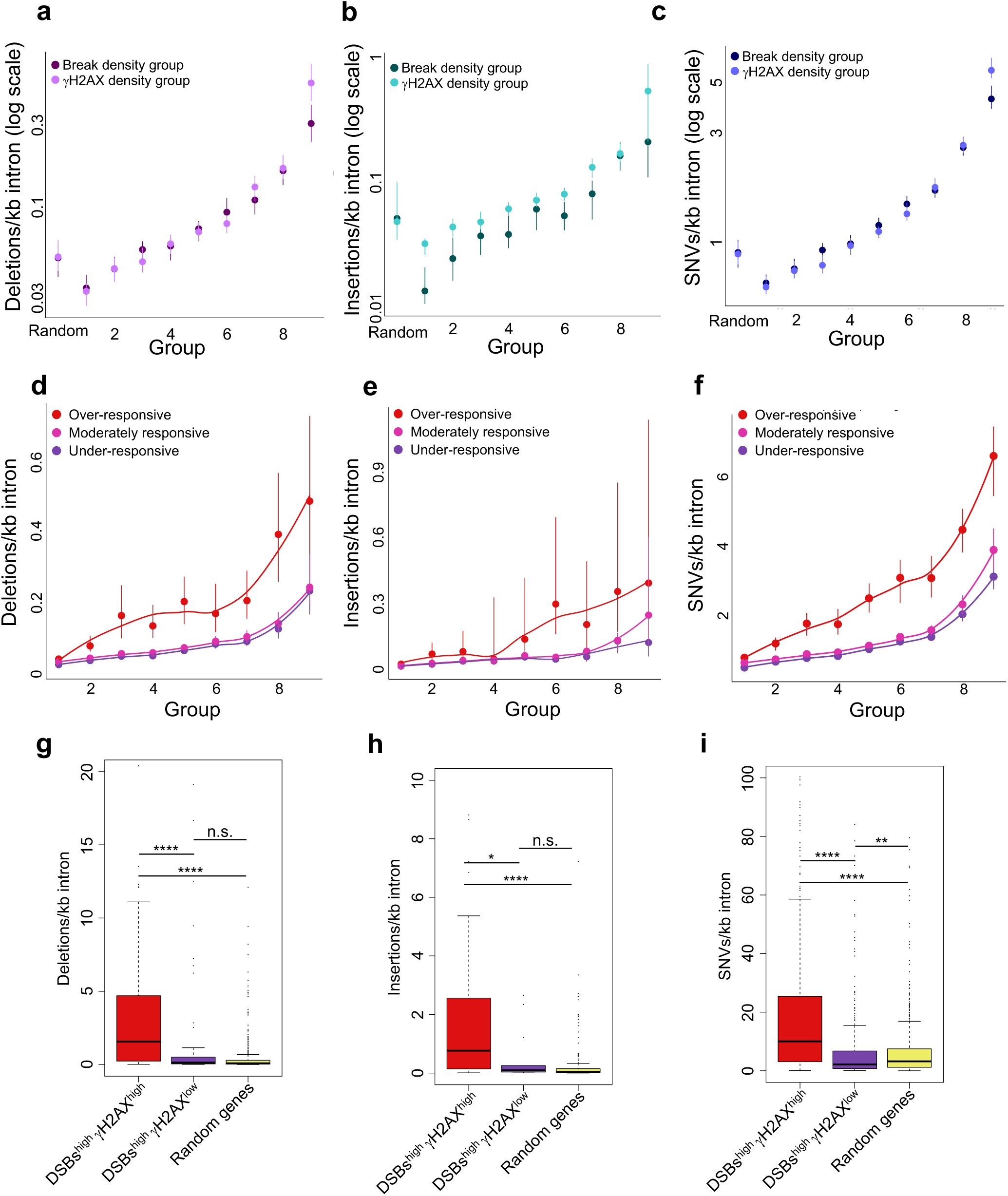
transcriptional DSBs with high turnover are especially susceptible to mutations. **a-c**. Intronic deletion density (**a**), insertion density (**b**), and SNV density (**c**) for genes categorized by their break or γH2AX density (excluding zero break genes). Random genes are included for reference. **d**-**f**, Rate of intronic mutations across genes categorized by their break density and further divided into three groups based on their γH2AX response to DSBs. **g-i**. Boxplot of intronic deletion density (**g**), insertion density (**h**), and SNV density (**i**) for DSBs^high^ γH2AX^high^ vs. DSBs^high^ γH2AX^low^ vs. randomly selected genes.

To disentangle the effects of DSB density from repair and turnover, we modelled the relationship between γH2AX density and break density using nonlinear least squares (NLS) regression. The residuals from this model were then used to classify genes in each group into three subgroups: Under-responsive genes (negative residuals), moderately responsive genes (residuals near zero), and over-responsive genes (positive residuals). Notably, over-responsive genes showed the most increase in mutation density with increased DSBs (Fig. 8D-F). This suggests that excessive γH2AX accumulation at DSB sites, which reflects recurrent DSB formation and turnover, and heightened repair activity, is error-prone. Furthermore, DSBs^high^ γH2AX^high^ genes, but not DSBs^high^ γH2AX^low^ genes, demonstrated intronic mutation enrichment, compared to randomly selected genes (Fig. 8G-I). Collectively, these results suggest that transcription stress sites are prone to mutations, likely due to a combination of high DSB turnover, clustering, and defective repair. These findings have important implications for understanding how transcription-associated genome instability contributes to cancer evolution, as recurrent DSB formation and error-prone repair may drive the accumulation of oncogenic mutations over time.

## Discussion

Cellular processes–particularly those upregulated in cancer such as replication and transcription–play a significant role in genomic instability^54^. Replication-dependent instability, caused by replication stress^4,55^, has been extensively studied, revealing its landscape^56,57^, causes, consequences^58^, and potential for therapeutic targeting^59,60^. Transcription-replication-dependent genomic instability^61^, particularly transcription-replication conflicts (TRCs)^62^, has also been well characterized, with studies examining its genomic distribution, influencing factors, and the impact of transcription-replication orientation^63^. Recently, attention has shifted toward transcription-dependent, replication-independent genomic instability, namely, transcriptional DSBs. While research has begun to uncover their causes and implications in disease^17,64^, several key aspects remain underexplored, including genomic hotspot patterns, repair mechanisms, and their impact on cancer evolution. By utilizing multi-omic approaches, we delineated the landscape of transcription stress and revealed that SEs shape the transcriptional program and contribute to the formation of genomic instability hotspots in cancer.

SEs determine cell identity in normal cells^31^. In cancer cells, SEs enhance the transcription of oncogenes to promote cell growth and proliferation ^6,8,65,66^. Previous studies have established that highly transcribed genes are prone to transcription-associated genome instability^13,15,20,30,67^. However, our data reveal that this relationship is significantly amplified in SE-regulated genes, suggesting that transcriptional stress is not merely a function of high gene expression but is modulated by SE activity. The enrichment of highly expressed SE-regulated genes among transcription stress sites was over 30-fold higher than expected by chance, far exceeding that of highly expressed TE-regulated genes or highly expressed genes lacking enhancer regulation. This specificity suggests that SEs impose a unique transcriptional burden that predisposes their target genes to DNA damage. The functional implications are significant, as many SE-regulated genes encode critical oncogenes, including CCND1, NEAT1, miR21, and MYC, which play essential roles in tumorigenesis.

The pronounced enrichment of DSBs at SE-regulated oncogenes was considerably higher in cancer cell lines compared to their normal counterparts, indicating that SEs primarily drive genomic instability in cancer cells. SEs have previously been recognized as hyper-mutated regions that can alter the expression of target genes and contribute to carcinogenesis^6^. Our findings advance this understanding by demonstrating that SEs also dictate mutational susceptibility at their target genes by causing hyper-transcription.

Although sBLISS captures a snapshot of the DSB landscape at the time of fixation, we could estimate the repair and turnover of DSBs across genes, revealing how transcription stress sites particularly undergo high DSB turnover and repair efficiency. To accommodate this, these regions experience constant activation of the DNA damage response (DDR) and H2AX phosphorylation. Interestingly, not all DSB-enriched genes exhibit such efficient repair signaling, suggesting two distinct types of DSB-bearing genes in the genome: genes that undergo frequent breakage-repair cycles and are actively recognized by the repair machinery, and genes that, despite being DSB-rich, experience infrequent breakage-repair cycles and remain persistent. These persistent DSBs, found in regions with low expression, have a lower demand for repair. We speculate that these latter genes experience breakage in a more programmed manner compared to genes with high turnover, which break as a byproduct of transcription. The nature of these DSBs requires further investigation.

Additionally, we demonstrate that genes with efficient DSB repair signaling are more vulnerable to mutations than genes with comparable DSB levels but less effective repair signaling, emphasizing the importance of the repair process itself in mutagenesis at transcription stress sites.

In summary, our findings highlight SEs as critical determinants of transcriptional stress and genomic instability in cancer. By integrating multi-omic data, we demonstrate that SE-driven hyper-transcription uniquely predisposes oncogenes to DNA breakage and mutation, revealing a previously underappreciated mechanism by which SEs shape the cancer genome. These insights not only deepen our understanding of transcription-dependent DNA damage but also point to SE-associated transcription stress as a potential vulnerability that could be therapeutically exploited. Further investigation into the distinct dynamics of DSB repair at these loci may uncover new strategies to modulate genome stability in cancer.

## Method details

### Cell culture and experimental conditions

MCF7 (HTB-22) and T-47D (HTB-133) cells were grown in RPMI supplemented with 10% (vol/vol) fetal bovine serum FBS (GIBCO), glutamine, and penicillin/streptomycin. HEK-293 (CRL-1573), MDA-MB-468 (HTB-132), MDA-MB-231 (HTB-26), and HeLa cells (CCL-2) were grown in DMEM supplemented with 10% (vol/vol) FBS, glutamine, and penicillin/streptomycin. A-4098 cells (HTB-44) were grown in. EMEM supplemented with 10% FBS, glutamine, and penicillin/streptomycin. SH-SY5Y (CRL-2266) were grown in DMEM/F12 supplemented with 10% FBS, glutamine, and penicillin/streptomycin. HMLE cells were grown in Promocell mammary epithelial cell basal media (C-21010) with added supplements (c-93110), whereas MCF10A were grown on DMEM/F12 supplemented with 5% Horse serum, 20 ng/ml EGF, 0.5 mg/ml Hydrocortisone, 100 ng/ml Cholera toxin, 10 mg/ml Insulin and Pen/Strep. MCF-7 cells were synchronized in the G1 cell cycle stage by culturing in serum-free media for 72 hours followed by replacing the media with 5% FBS media for 5 hours. Cells were grown at 37°C under a humidified atmosphere with 5% CO2. Cells were routinely authenticated by STR profiling, and tested for mycoplasma, and cell aliquots from early passages were used.

Cells were treated with DMSO (Sigma-Aldrich), dBET6 (Sigma-Aldrich, SML2638; 100 nM), or CDK9i (NVP-2) (MCE, Cat# Y-12214A; 100 nM) for 4 hours. Cells were treated with DMSO or DNA-PKci (NU-7441, Selleckchem; 2 μM) for 5 hours.

### Transcription stress score

Our sBLISS, DRIP-seq, TOP1cc, and TOP1 ChIP-seq data used for transcription stress score calculation are already published^13^ and are available at GEO: GSE241309. For each dataset, the signal density per gene was first converted to an empirical p-value based on its rank within the distribution of all genes. Higher signal values correspond to lower p-values. These p-values were then transformed into Z-scores using the standard normal inverse cumulative distribution function:

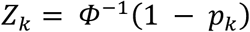

where *pₖ* is the empirical p-value for dataset *k*, and *Φ⁻¹* denotes the inverse of the standard normal cumulative distribution function.

The individual Z-scores from all four datasets were combined per gene using the Liptak weighted Z-test (unweighted version):

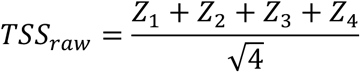

For better representation, the combined score was further min-max normalized and then stretched by a power transformation to emphasize differences:

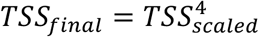

Genes not among the top 1000 in one or more parameters were excluded.

### Identification of Transcription Stress-Associated Domains using Multivariate HMM

To identify genomic regions simultaneously enriched for markers of transcription stress, we applied a multivariate hidden Markov model (HMM) using the *mhsmm* R package. The hg38 genome was divided into 1 kb non-overlapping bins, and mean signals for TOP1, TOP1cc, R-loops, and DSBs were calculated per bin. Signal intensities were capped at the 0.5th and 99.5th percentiles to minimize the influence of outliers. A two-state multivariate HMM was initialized with uniform state priors and a transition matrix favouring self-transition (0.95 probability). Emissions were modelled using multivariate Gaussian distributions with means offset by ±0.5 standard deviations and shared covariance. The model was trained by maximum likelihood estimation, and Viterbi decoding was used to assign each bin to a state. The state with the highest average signal across all datasets was designated as the “co-marked” state. To refine the identified regions, we selected bins showing high signal (above the 95th percentile) in at least three of the four datasets, merged adjacent bins separated by ≤1 kb, and filtered to retain regions ≥3 kb. ^13^

### Gene sets compilation and enrichment analysis

Gene sets tested for enrichment in this manuscript were defined as follows: SE-regulated genes for each cell line were downloaded from SEdb2.0^31^. Samples IDs used were: Sample_01_0046 for MCF7, Sample_02_1273 for HeLa, Sample_00_0015 for IMR90, Sample_02_0434 for HEK-293, Sample_02_0097 for T-47D, Sample_02_0095 for MDA-MB-468, Sample_02_0219 for MDA-MB-231, Sample_02_0643 for MDA-MB-436, Sample_02_1812 for SHSY5Y, Sample_02_0056 for HMLE, Sample_02_0268 for A4098. A list of annotated oncogenes was downloaded from OncoKB^68^. Highly expressed genes are defined as the top 1000 expressed genes. TE-regulated genes are defined as genes that overlap enhancer regions and are not annotated by an SE. Non-E genes are genes that do not overlap any enhancer element. enrichment was calculated by dividing the number of observed overlapping genes by the theoretical number of genes predicted under a random model. One-tailed (upper-tailed for values > 1, and lower-tailed for values < 1) hypergeometric test using the phyper function in R to determine the statistical significance of the observed enrichment.

### In-suspension break labelling *in situ* and sequencing (sBLISS)

sBLISS was conducted as previously described^41,69^. In summary, 10^6^ cells were fixed in 2% paraformaldehyde in 10% FCS/PBS for 10 minutes at room temperature. The fixation was quenched with 125 mM glycine for 5 minutes at room temperature, followed by another 5 minutes on ice and two washes in ice-cold PBS. Cells were lysed for 60 minutes on ice, and their nuclei were permeabilized for 60 minutes at 37°C. Next, nuclei were rinsed twice with CutSmart Buffer containing 0.1% Triton X-100 (CS/TX100), and double-strand break (DSB) ends were blunted in situ using NEB’s Quick Blunting Kit for 60 minutes at room temperature. The blunted nuclei were then washed twice with 1x CS/TX100 before in situ ligation of the sBLISS adapters to the DSB ends. Adaptor ligation was carried out with T4 DNA Ligase for 20-24 hours at 16°C, with BSA and ATP added. Following ligation, the nuclei underwent two washes with 1x CS/TX100, and genomic DNA was extracted using Proteinase K at 55°C for 14-18 hours while shaking at 800 rpm. Proteinase K was then heat-inactivated for 10 minutes at 95°C, followed by extraction using Phenol:Chloroform:Isoamyl Alcohol, Chloroform, and ethanol precipitation. The purified DNA was sonicated in 100 μL of ultra-pure water using Covaris M220 for 60 seconds. Sonicated samples were concentrated with AMPure XP beads (Beckman Coulter), and fragment sizes were evaluated using a BioAnalyzer 2100 (Agilent Technologies), targeting a range of 300 bp to 800 bp with a peak around 400-600 bp. The sonicated DNA was then in vitro transcribed using the MEGAscript T7 Kit for 14 hours at 37°C. After RNA purification and ligation of the 3’-Illumina adaptors, the RNA underwent reverse transcription. The final library indexing and amplification step was performed with NEBNext® Ultra™ II Q5® Master Mix.

sBLISS fastq files are initially de-multiplexed using sample barcodes. Quality control is performed with trim galore to eliminate residual adapters, trim reads to a base quality of at least 20, and filter out short reads smaller than 20 bp. Initial and final sample qualities are assessed with fastqc. Quality-processed fastq files are aligned to the GRCh38 assembly with hisat2, then sorted and indexed using samtools. The resulting bam files are de-duplicated with umi-tools, utilizing genomic coordinates and Unique Molecular Identifiers (UMI). Custom Python and R scripts are employed to identify read start positions and convert bam files to bigwig format for downstream analysis. An additional custom R script is used to discard a blacklist of positions, primarily within centromeres.

### RNA-seq and GRO-seq data processing

Bulk RNA-seq processed data for untreated MCF7, HeLa, and T-47D cell lines were obtained from ENCODE^70,71^, while data for MDA-MB-436 and MDA-MB-468 were downloaded from GEO (GSE212143)^72^. For MCF7, the following datasets were used: ENCFF967AOT, ENCFF328CKZ, ENCFF921PJP, ENCFF930TQB, ENCFF030IUO, ENCFF878CJF, ENCFF057ITQ, ENCFF317XCR, ENCFF985KJE, and ENCFF456OYZ. For HeLa, we used ENCFF750TQI, ENCFF622TLZ, ENCFF689BFU, ENCFF151QTZ, ENCFF262EJQ, ENCFF878CJF, ENCFF964OAC, ENCFF346JDS, and ENCFF638MLA. For T-47D, we used ENCFF234TQB, ENCFF412DJX, ENCFF475SHD, and ENCFF846PLI. TPM values were averaged across replicates for each cell line to determine the expression rank of each gene.

RNA-seq data for MCF7 after CDK9i treatment was downloaded from <u>GSE129012</u> (4h timepoint). And GRO-seq data for MCF7 after E2 treatment was downloaded from GSE27463 (40 min timepoint).

### RT-qPCR and Spike-in RT-qPCR

Total RNA was isolated utilizing TRI reagent (Biolab), following the manufacturer’s guidelines for the phenol/chloroform extraction technique. cDNA was synthesized from 1 μg of RNA using the QScript cDNA synthesis kit (Quantabio). The SYBR Green PCR Master Mix (Applied Biosystems) was employed for qRT-PCR. All assays were conducted in triplicate. Experiments were performed to assess the levels of cDNA using primers targeting: Chicken RPL4 (FW, GAGTGACTACAACCTGCCGA; REV, TTGGCGTATGGGTTCAGCTT), SIK1 (FW, CTCCGGGTGGGTTTTTACGAC; REV, CTGCGTTTTGGTGACTCGATG), ID3 (FW, GAGAGGCACTCAGCTTAGCC; REV, TCCTTTTGTCGTTGGAGATGAC), XBP1 (FW, CCCTCCAGAACATCTCCCCAT; REV, ACATGACTGGGTCCAAGTTGT), CCND1 (FW, GCTGCGAAGTGGAAACCATC; REV, CCTCCTTCTGCACACATTTGAA), HES1 (FW, CCTGTCATCCCCGTCTACAC; REV, CACATGGAGTCCGCCGTAA), FOS (FW, TCCCATCGGTCCACTAGGTTT; REV, AGGGCTGCACTGAGTTCTTTG), JUNB (FW, ACGACTCATACACAGCTACGG; REV, GCTCGGTTTCAGGAGTTTGTAGT), TFF1 (FW, CCCTCCCAGTGTGCAAATAAG; REV, GAACGGTGTCGTCGAAACAG), UBC (FW, ATTTGGGTCGCGGTTCTTG; REV, TGCCTTGACATTCTCGATGGT), HPRT (FW, TGACACTGGCAAAACAATGCA; REV, GGTCCTTTTCACCAGCAAGCT), MYC (FW, ATGCCCCTCAACGTGAACTTC; REV, CGCAACATAGGATGGAGAGCA), XRCC4 (FW, ATGTTGGTGAACTGAGAAAAGC; REV, GCAATGGTGTCCAAGCAATAAC), GREB1 (FW, TGGTCCGTAATGCACAAGGG; REV, CTGCGTTTAGTGAGGGGTGA), GATA3 (FW, GCCTCTGCTTCATGGATCCC; REV, CACACTCCCTGCCTTCTGTG), ACTB (FW, TTTTGGCTATACCCTACTGGCA; REV, CTGCACAGTCGTCAGCATATC).

In RT-qPCR experiments, Ct values were normalized to HPRT levels. For spike-in RT-qPCR, chicken RNA was added in proportion to the number of cells in each sample, and Ct values were normalized to chicken RPL4.

### γH2AX ChIP-seq

MCF7 cells (∼10^6^) were crosslinked with 1% formaldehyde (methanol free, Thermo Scientific 28906) for 10 min at room temperature and quenched with glycine, 125 mM final concentration. Fixed cells were washed twice in PBS and incubated in sonication buffer (0.5M NaCl, 0.5%SDS RIPA buffer containing PMSF, protease and phosphatase inhibitors) for 30min on ice. Cells were sonicated using Covaris M220 for 10min to produce chromatin fragments of ∼200-300 bp. The sheared chromatin was centrifuged 10min at maximum speed. From the supernatant, 50ul were saved as input DNA and the rest was diluted in RIPA buffer without NaCl and SDS. The chromatin was immunoprecipitated by incubation with 5 μg of gamma-H2AX (phospho S139; Abcam, AB2893) antibody pre mixed with 100ul Dynabeads Protein G (ThermoFisher, 10004D) and incubated over-night at 4°C with rotation. Immunoprecipitates were washed twice with RIPA buffer + 150mM NaCl, twice with RIPA buffer + 300 mM NaCl, and twice with TE buffer. The chromatin was eluted from the beads with 200ul of direct elution buffer (10mM Tris pH 8, 0.3M NaCl, 5mM EDTA, 0.5% SDS) and incubated overnight at 65°C to reverse the cross-linking. Samples were treated with RNAse for 1h at 37 °C and with proteinase K at 55 °C for 2 h. DNA was cleaned up by QIAquick PCR purification column (Qiagen), according to the manufacturer’s instructions, and eluted in 30 μl of elution buffer. The ChIPed and the Input DNA were used for real-time PCR or to prepare libraries by Truseq (Ilumina) and sequenced in Nextseq (Illumina). ChIPed and input fastq files are quality controlled with trim galore to remove residual adapters, trim reads to base quality of at least 20 and filter out short reads with size smaller than 20 bp. Initial and final sample qualities are evaluated with fastqc. The quality processed fastq files are aligned to GRCh38 genome with hisat2 followed by sorting and indexing with samtools. The utility bamCompare of deepTools is applied on the bam files to create a peak bigwig file. This operation is set to ignore duplicates, discard a blacklist of regions, normalize by RPKM, and compare by ratio in tiles of 50 bp. γH2AX ChIP-seq data for HeLa and IMR90 were downloaded from GEO: GSM4564685 and GEO: GSM2067927, respectively. γH2AX density per gene was computed using bigWigAverageOverBed, which calculates the average signal from the γH2AX bigWig track over annotated gene intervals.

### Statistical modelling and residual-based classification

For multivariate logistic regression, genes were categorized based on whether they overlapped transcription stress sites identified by Hidden Markov Modeling (HMM). Logistic regression was performed using the glm function in R (family = binomial).

Multivariate Least Absolute Shrinkage and Selection Operator (LASSO) regression was conducted using the glmnet package in R, with 10-fold cross-validation (cv.glmnet) to select the optimal regularization parameter (λ). The model was trained to classify genes based on transcription stress site overlap or DSB enrichment. Odds ratios were derived from the final model coefficients to assess the relative predictive power of each feature.

Generalized additive model (GAM) was applied to model the relationship between each variable and γH2AX across all genes in (Fig. 5A), and for genes classified into their overlapping chromatin states in (Fig. 5H). The model was constructed using the gam function in the mgcv package in R.

To dissect the effects of absolute DSBs from DSB turnover on mutation frequency, we modelled the relationship between break density and transcription stress by nonlinear least squares (NLS) regression using the following equation:

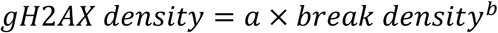

The model was fitted using the nls function in R, with initial parameter estimates for “a” and “b” set to 0.1. After fitting the model, residuals were extracted and used to classify genes into three response categories based on predefined thresholds: genes with residuals > 0 were classified as ‘Over-responsive,’ genes with residuals < 0 as ‘Under-responsive’ and genes between –0.3 and 0.3 as ‘Moderately responsive’.

### Classifying individual DSB sites based on γH2AX

Each break site was assigned a γH2AX coverage level (γH2AX rank). To ensure a comparable distribution of break scores (i.e., the number of DSBs at the same site) between the two groups, break sites were classified into four categories: those with a break score of 1 (the large majority), those with a break score between 2 and 4, those with a break score between 5 and 10, and those with a break score between 11 and 50. Within each of the four categories, 1% of the top and bottom γH2AX-ranked breaks are selected. All categories are then re-combined so that the top set of breaks and the bottom set of breaks exhibit a similar break score distribution. Mapping to chromatin states is then computed and plotted for these sets.

### Chromatin states

Chromatin states bed files were downloaded from ENCODE^70,71^. For MCF7, ENCFF631HCR file was used. For HeLa, ENCFF555HXM file was used.

### Transient transfection

Transient transfection of siXRCC4 (Dharmacon; SMARTpool siRNA, Catalog ID: L-004494-00-0005) and siSc (Dharmacon; D-00181010) was carried out using Lipofectamine (ThermoFisher). On the day of transfection, cells were seeded to achieve a confluency of 60-70%. The transfection solution was prepared by mixing 1.5 ml of serum and antibiotic-free RPMI with 30 µL of Lipofectamine and incubating for 5 minutes before adding a mixture of 1.5 ml RPMI and 0.6 nmoles of siRNA, followed by a 20-minute incubation at room temperature. This mixture was then added to cells in a 10 cm plate cultured in antibiotic and serum-free media. The media was replaced with fully supplemented media after 5-6 hours, and the cells were incubated in the incubator for 48 hours.

### HiC data analysis

Normalized Hi-C contact matrices for MCF7 and MCF10A cells were obtained from GEO (GSE66733).

Genomic bins were intersected with all genes, and each bin was assigned to a single gene. Bins overlapping multiple genes were excluded from the analysis.

In inter– and intra-group metrics calculation, genes were stratified into nine groups based on their break density, excluding genes with zero breaks. Intra-group and inter-group interaction scores were computed as follows: Intra-group interactions: For each gene, interaction scores with all other genes within the same group were summed and normalized by the total number of genes in that group. Inter-group interactions: For each gene, interaction scores with all genes outside its group were summed and normalized by the total number of genes in all other groups. Genes, along with their interaction scores, were ordered based on break density. The data was visualized using loess smoothing.

### COSMIC mutations data analysis

The file Cosmic_NonCodingVariants_Tsv_v101_GRCh37.tar, containing all noncoding variants, was downloaded from COSMIC. This file was first filtered for breast cancer samples and separated based on the type of mutation into SNVs, deletions, and insertions mutation files. These files were then converted into genomic ranges objects and filtered to contain only mutations within introns. To calculate mutation density, mutations per gene were counted and normalized by the sum of intron length.

## Acknowledgments

We sincerely thank the members of the Aqeilan lab for their valuable discussions and insights. We are deeply grateful to Prof. Sheera Adar and Dr. Yotam Drier for their thoughtful guidance and support as members of the PhD committee. We also appreciate the invaluable assistance of Dr. Abed Nasereddin and Dr. Idit Shiff from the Core Research Facility of the Hebrew University-Hadassah Medical School. Additionally, we extend our sincere gratitude to Prof. Yinon Ben Neriah for generously providing dBET6 and CDK9i. This study was supported by a grant from the Israel Science Foundation (ISF) [No. 1056/21]. We also acknowledge the support of the Carole and Andrew Harper Diversity Scholarship Program to O.H. and D.S, Science Training Encouraging Peace (STEP) Fellowship to D.S. and S.O. and the VATAT PhD scholarship to O.H.

## Author contributions

O.H. and R.I.A. conceived and designed the study. O.H. and J.M. performed computational analysis. S.O.F performed sBLISS on HMLE, MCF10A, MCF7, T47D, MDA-MB-436 and MDA-MB-468 cell lines. D.S. performed sBLISS on MCF7 cells synchronized in G1, SH-SY5Y cells, and NSCs. O.H. performed all other experiments. O.H. and R.I.A wrote the manuscript.

---

**Extended Data Figure 1.**
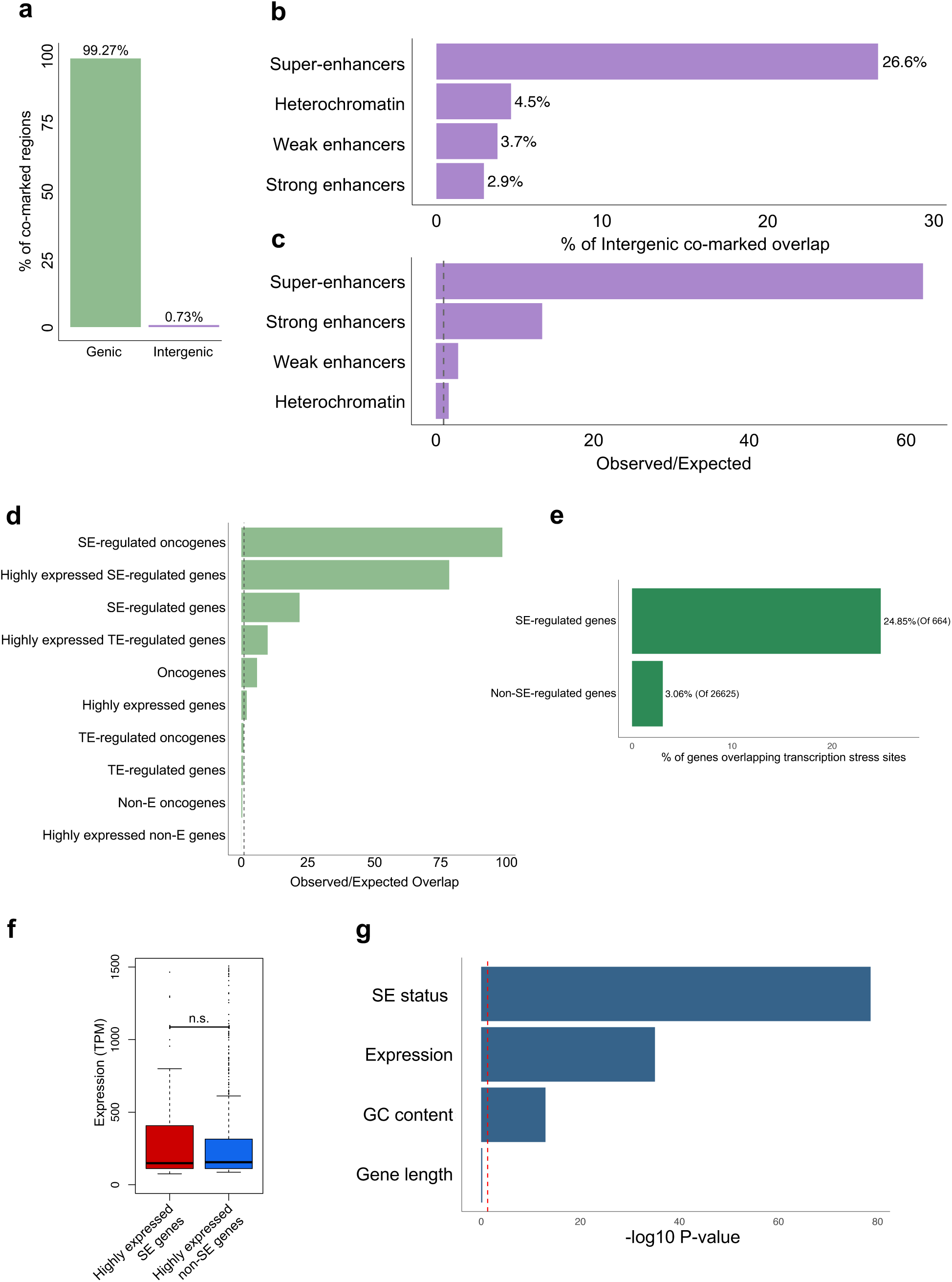
Characterization of transcription stress sites defined by HMM analysis. **a**, Distribution of transcription stress sites identified by HMM across genic and intergenic regions. Genic regions were defined as gene coordinates extended by ±10 kb flanking regions. **b,** Proportion of intergenic transcription stress sites identified by HMM overlapping key chromatin states and super-enhancers. **c,** Enrichment analysis showing observed over expected overlap of intergenic transcription stress sites with chromatin states and super-enhancers. **d,** Enrichment analysis of transcription stress sites within different gene categories. **e,** percentage of genes overlapping transcription stress sites as identified by HMM. **f,** Boxplot displaying expression levels of noted genes. **g,** Logistic regression analysis testing the predictive power of SE status, gene expression, GC content, and gene length for the presence of transcription stress sites within genes.

**Extended Data Fig. 2.**
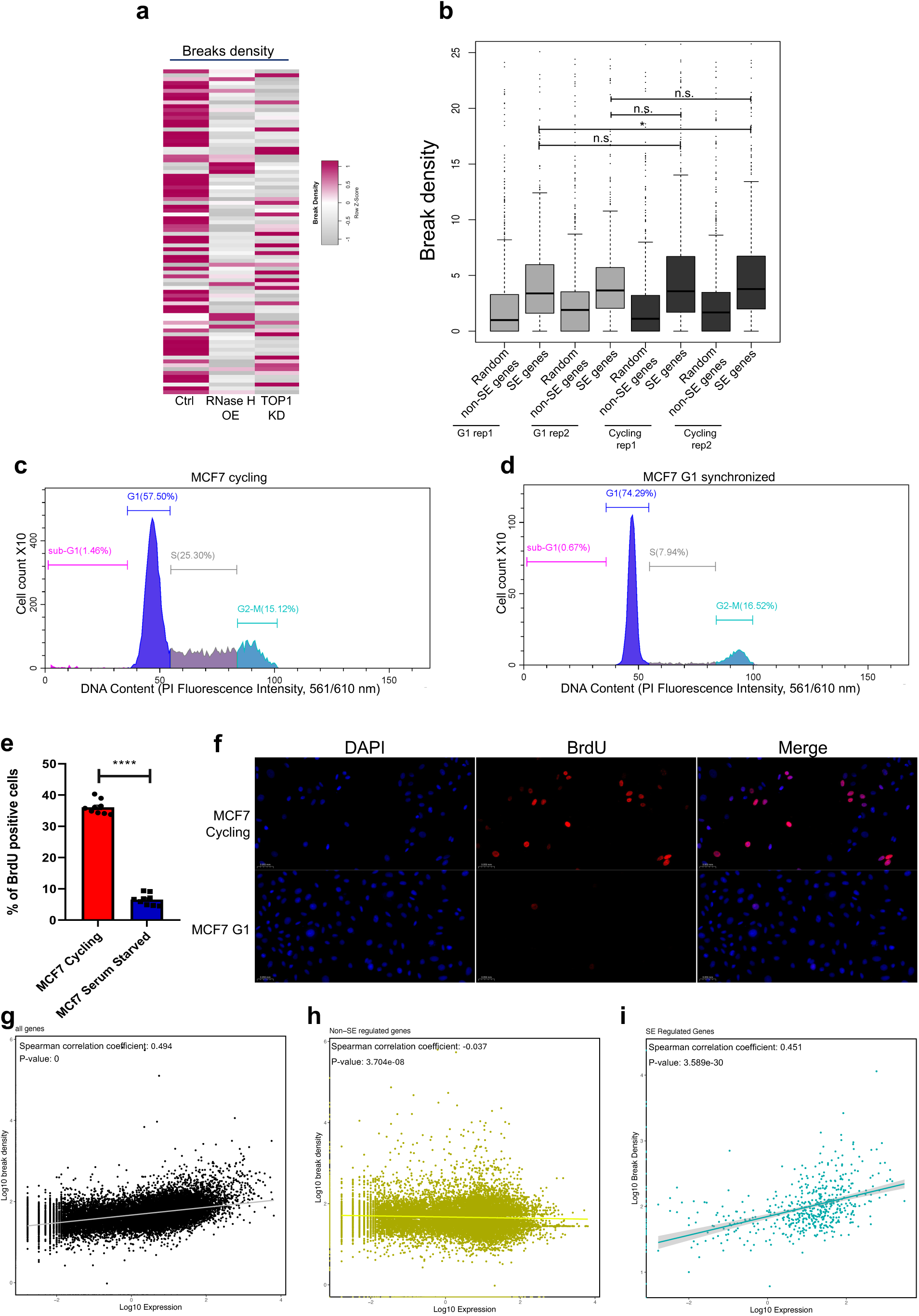
**a**, Heatmap illustrating break density of SE-regulated genes following RNase H overexpression and TOP1 KD. **b**, Boxplot showing break density of random genes or SE-regulated genes in MCF7 cells synchronized in G1 phase or cycling cells. **c-d,** Flow cytometry analysis of DNA content in MCF7 cells. (c) DNA content profile of asynchronously cycling MCF7 cells. (d) DNA content profile of MCF7 cells synchronized in G1 phase, demonstrating enrichment of the population in G1. Cells were stained with propidium iodide (PI) and analyzed by flow cytometry. **e**, percentage of BrdU positive cells in cycling vs. G1 synchronized MCF7 cells. **f**, representative images of BrdU incorporating cells. **g-I**, Scatter plot demonstrating the correlation between expression and break density for all genes (**g**), non-SE-regulated genes (**h**), and SE-regulated genes (**i**). Boxplot: center line indicates median; box limits represent upper and lower quartiles; whiskers extend to 1.5x interquartile range; dots indicate outliers. P-value was calculated using Mann Whitney U test. n.s. indicating p > 0.05, * indicating p < 0.05.

**Extended Data Fig. 3.**
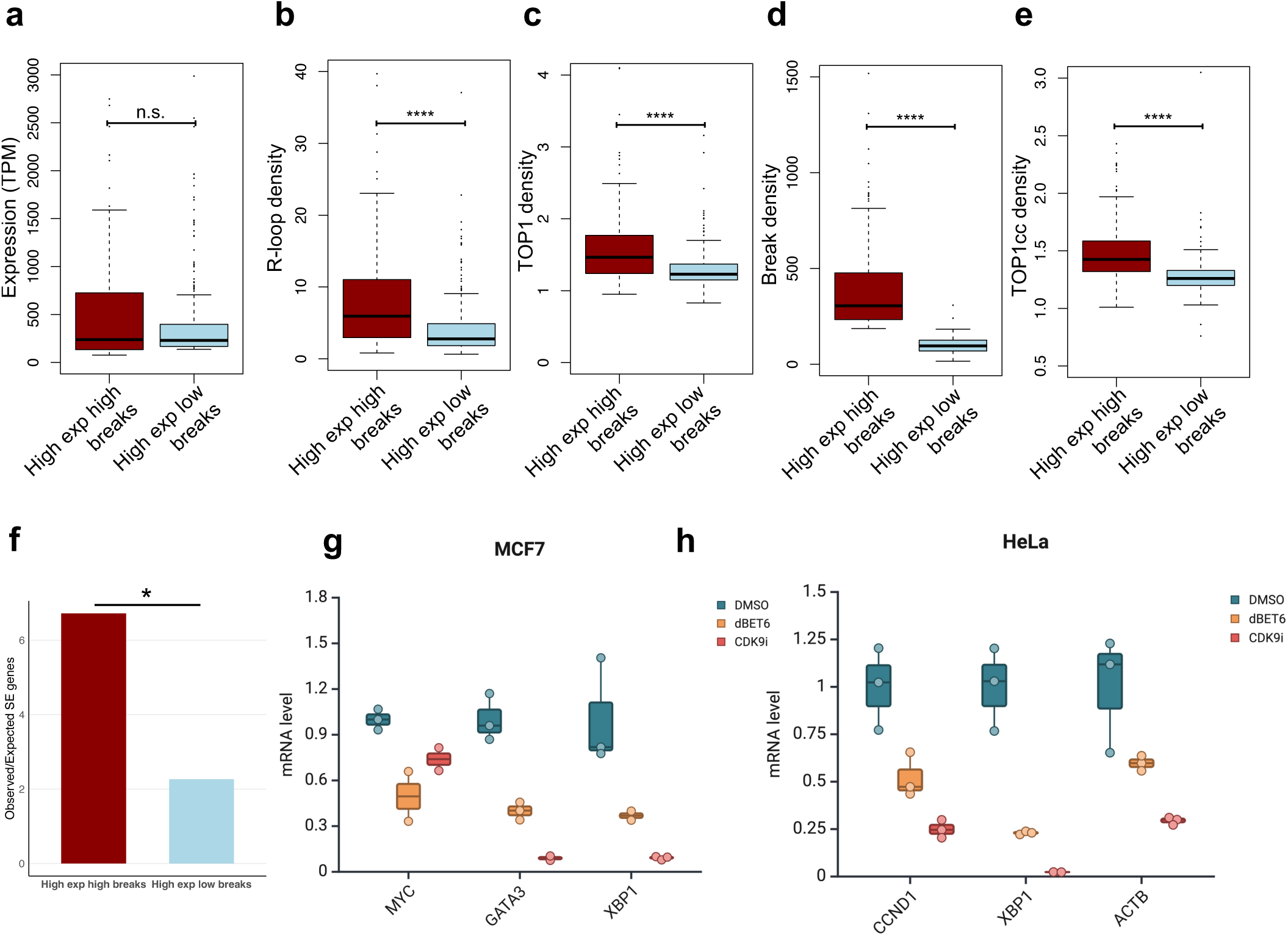
**a**-**e**, Boxplot showing expression levels (**a**), R-loop density (**b**), TOP1 density (**c**), break density (**d**), and TOP1cc density (**e**) for high expression high breaks genes versus high expression low breaks genes. **f**, observed to expected ratio of SE genes in high expression high breaks versus high expression low breaks genes. **g**-**h**, boxplot illustrating mRNA levels of sample SE-regulated genes in MCF7 (**g**) and HeLa (**h**) cells after dBET6 or CDK9i (100nM for 4 hours). Boxplot: center line indicates median; box limits represent upper and lower quartiles; whiskers extend to 1.5x interquartile range; dots indicate outliers. P-value was calculated using Mann Whitney U test. n.s. indicating p > 0.05, **** indicating p < 0.0001.

**Extended Data Fig. 4.**
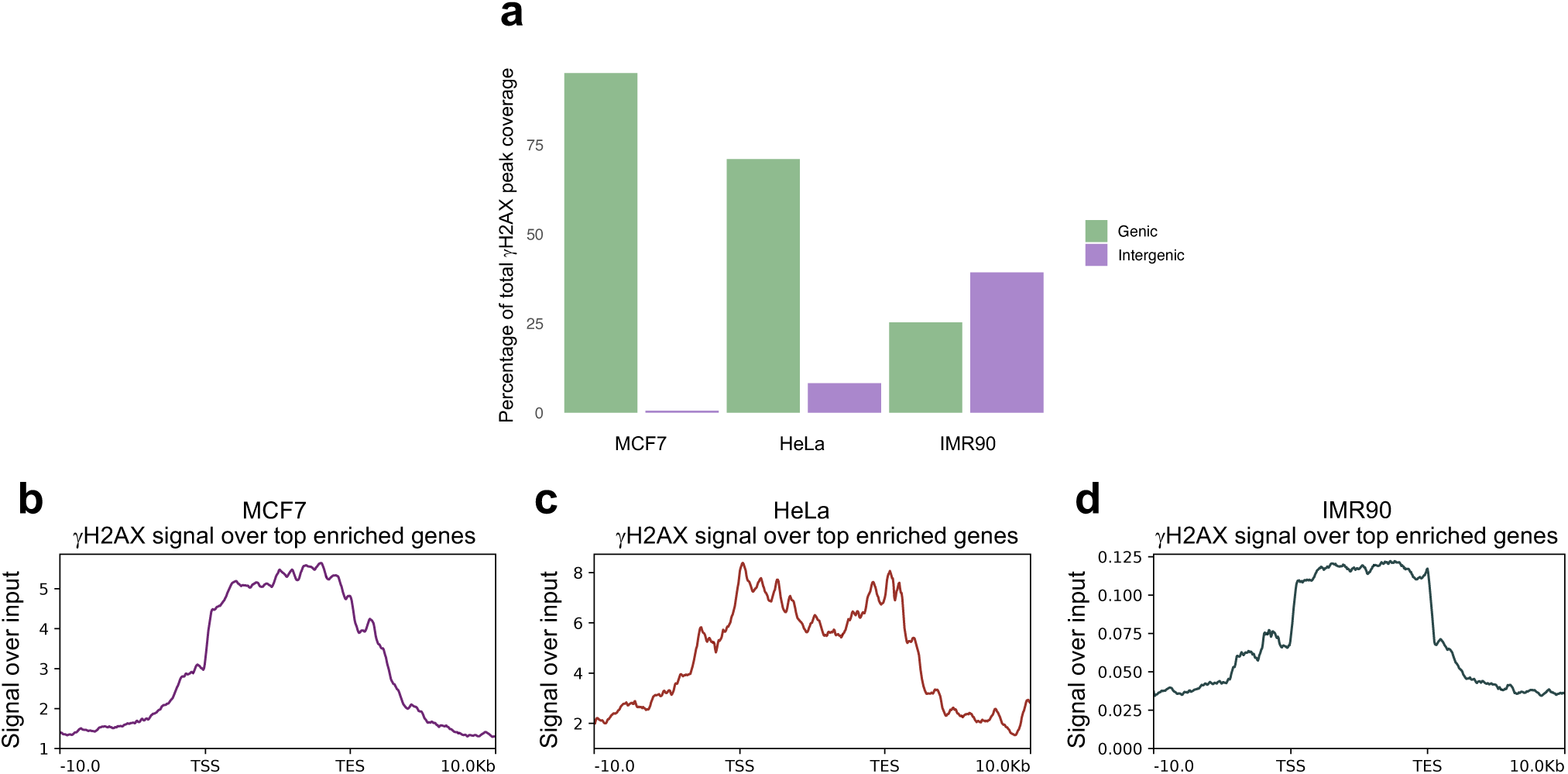
Distribution and enrichment of γH2AX signal at genic regions across different cell lines. **a**, Percentage of total γH2AX peak coverage occurring within genic and intergenic regions in MCF7, HeLa, and IMR90 cells. Genic regions were defined as gene coordinates extended by ±10 kb flanking regions. (**b**–**d**) γH2AX signal over input across the gene body and flanking regions (±10 kb) for the top enriched genes in (**b**) MCF7, (**c**) HeLa, and (**d**) IMR90 cells. TSS: transcription start site; TES: transcription end site.

**Extended Data Fig. 5.**
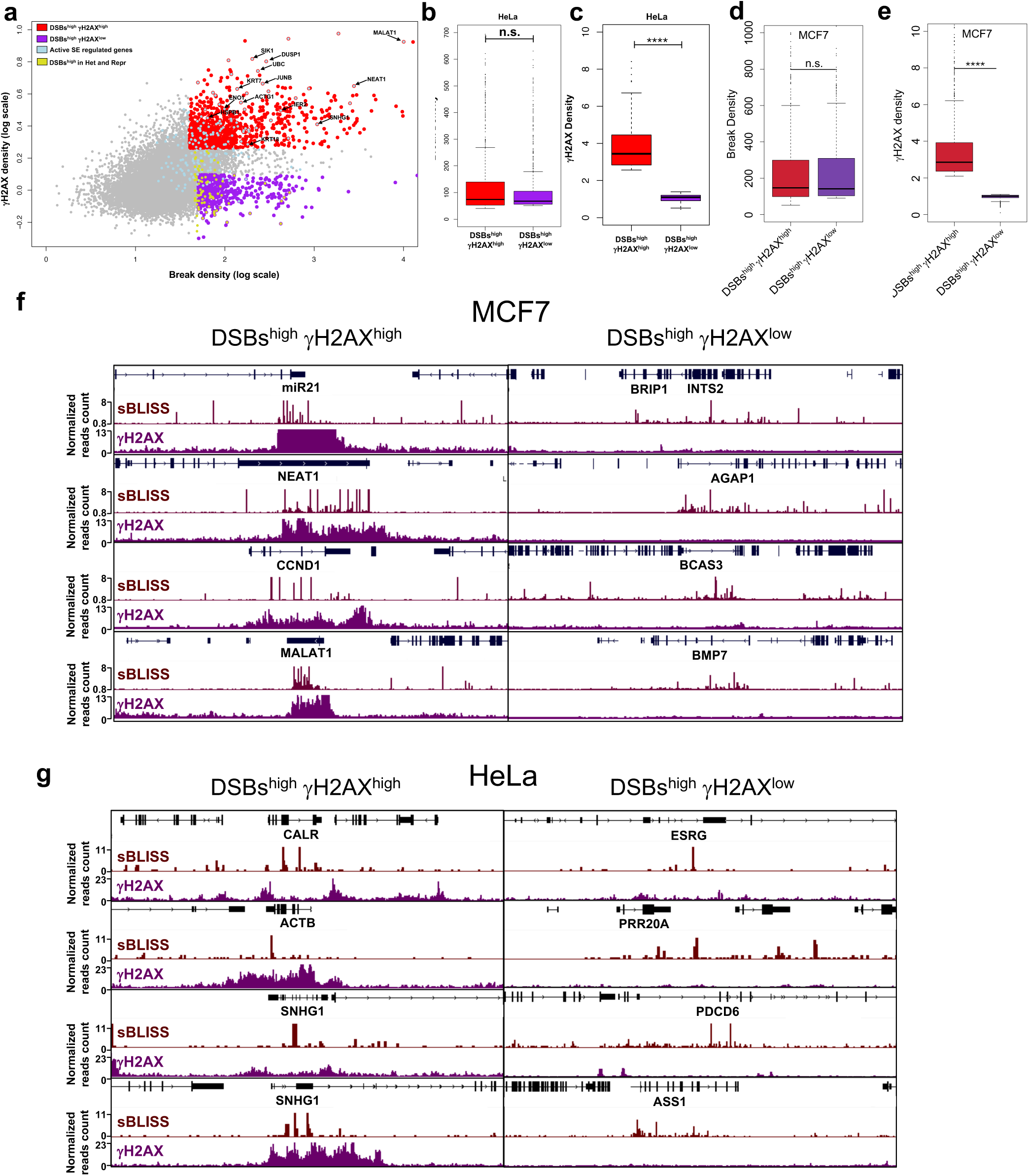
**a**, Scatter plot showing the correlation between break density and γH2AX density for each gene in HeLa cells. **b**-**e**, boxplots showing DSBs (b and d) and γH2AX density (c and e) for the two groups of genes defined in (a) and Figure 3F. **d-e,** Genome browser snapshots of representative examples of DSBs^high^ γH2AX^high^ and DSBs^high^ γH2AX^low^ genes in MCF7 (d), and HeLa (e) cell lines. Boxplot: center line indicates median; box limits represent upper and lower quartiles; whiskers extend to 1.5x interquartile range; dots indicate outliers. P-values were calculated using Mann Whitney U test. n.s. indicating p > 0.05, **** indicating p < 0.0001.

**Extended Data Fig. 6.**
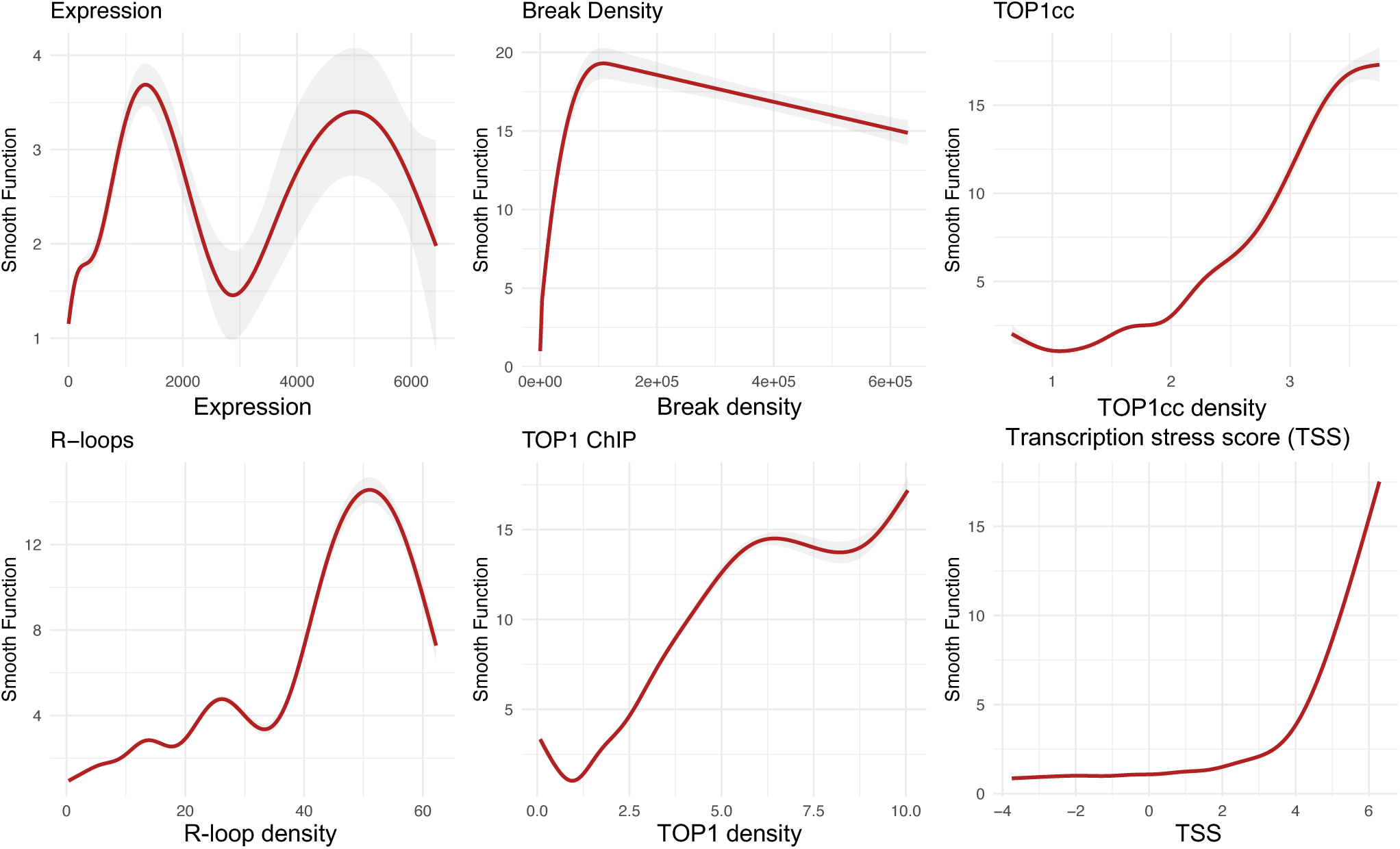
Smooth terms from Generalized Additive Models (GAMs) showing the effect of transcription-related predictors on γH2AX density. The y-axis shows the partial effect of each predictor (centered smooth function, s(x)) on γH2AX levels after accounting for non-linear relationships. The red line represents the fitted smooth function, and the shaded gray band indicates the 95% confidence interval.

**Extended data Fig. 7.**
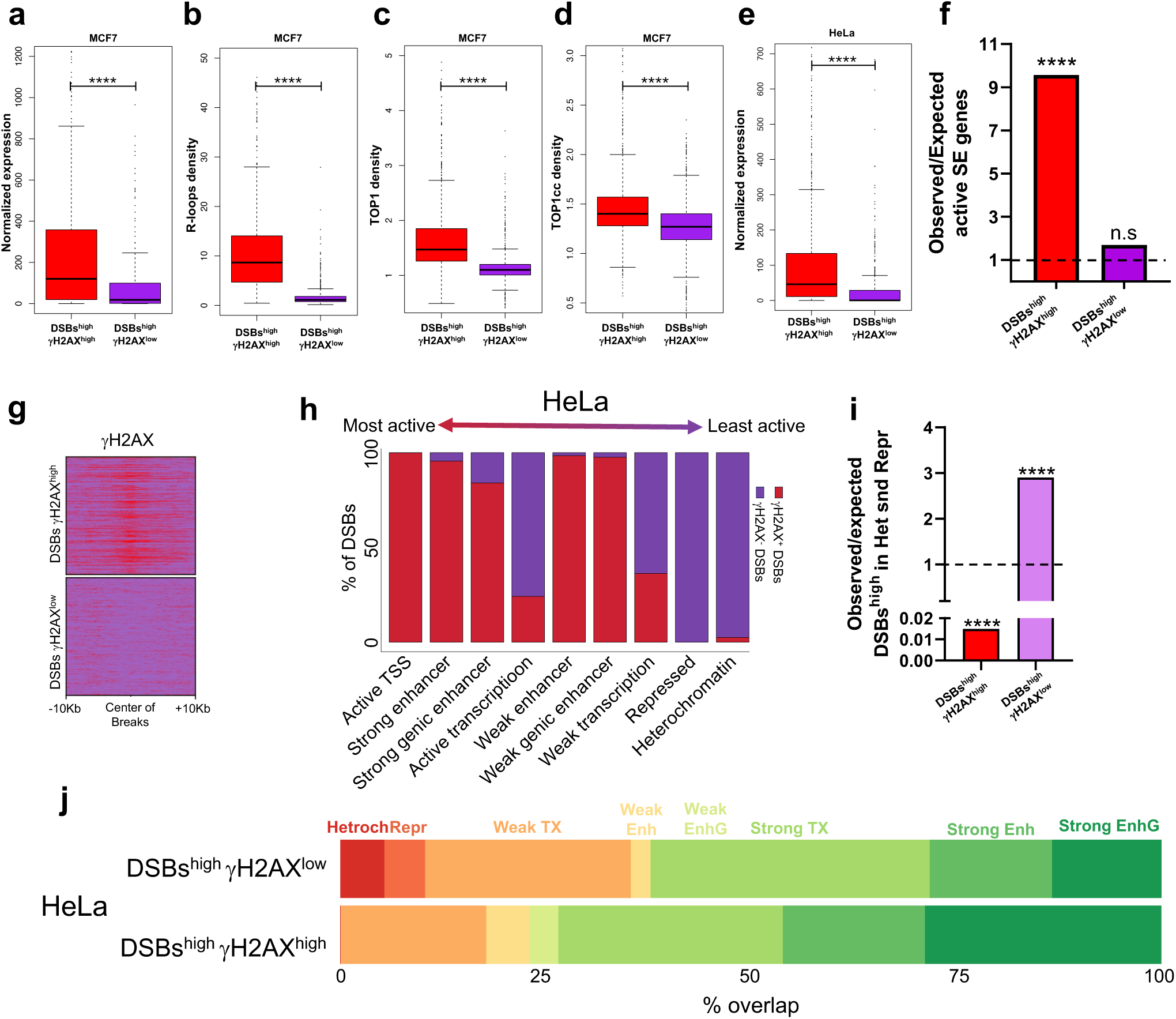
**a-d**, Boxplot showing expression levels (**a**), R-loop density (**b**), TOP1 density (**c**), TOP1cc density (**d**) for DSBs^high^ γH2AX^high^ and DSBs^high^ γH2AX^low^ genes in MCF7 cells. **e**, boxplot showing expression levels for DSBs^high^ γH2AX^high^ and DSBs^high^ γH2AX^low^ genes in HeLa cells. **f**, observed to expected ratio of active SE genes in DSBs^high^ γH2AX^high^ in HeLa cell line. **g**, heatmap showing the enrichment of γH2AX signal at the two defined sets of DSBs. **h**, The percentage of DSBs enriched with γH2AX (γH2AX^high^ DSBs) or lacking γH2AX (γH2AX^low^ DSBs) in each chromatin state of HeLa cell line. Chromatin states were ordered based on what is typically known about their transcriptional activity^73^. **i**, The ratio of observed to expected numbers of high DSB genes overlapping heterochromatin and repressed regions in both gene groups in HeLa cells. **j**, The percentage of overlaps between each gene group and chromatin states in HeLa. For each group, the number of overlaps was counted for each chromatin state and then divided by the total number of overlaps.

**Extended Data Fig. 8.**
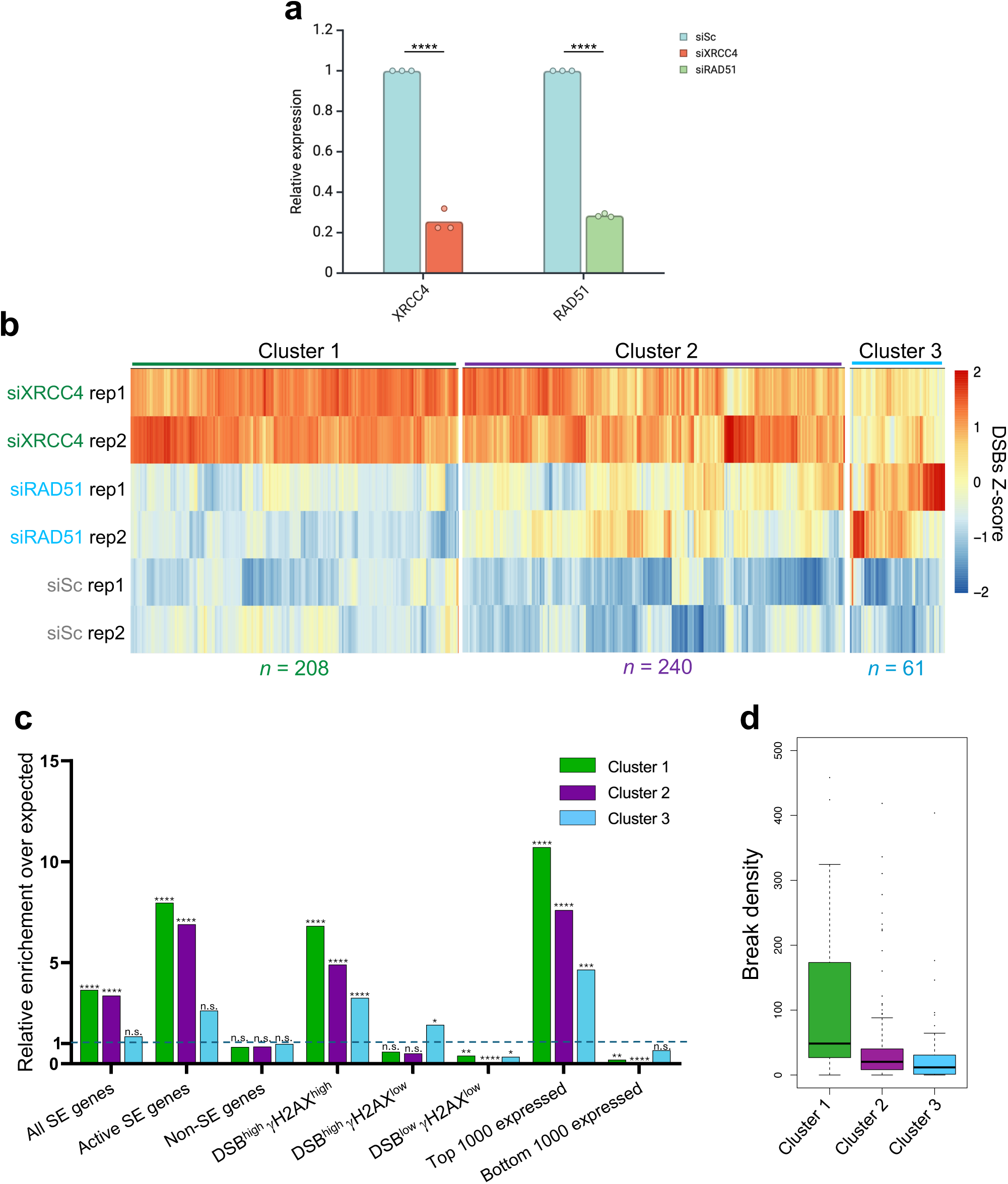
**a**, validation of XRCC4 and RAD51 knockdown via RT qPCR, each dot represents the mean of three technical replicates of a biological replicate. **b**, heatmap showing genes that significantly accumulated DSBs (p-value <0.05) upon XRCC4 or RAD51 KD. The genes were clustered by hierarchical clustering. c, enrichment of the three gene clusters in different gene sets. d, baseline break density of the three identified clusters.

**Extended data Fig. 9:**
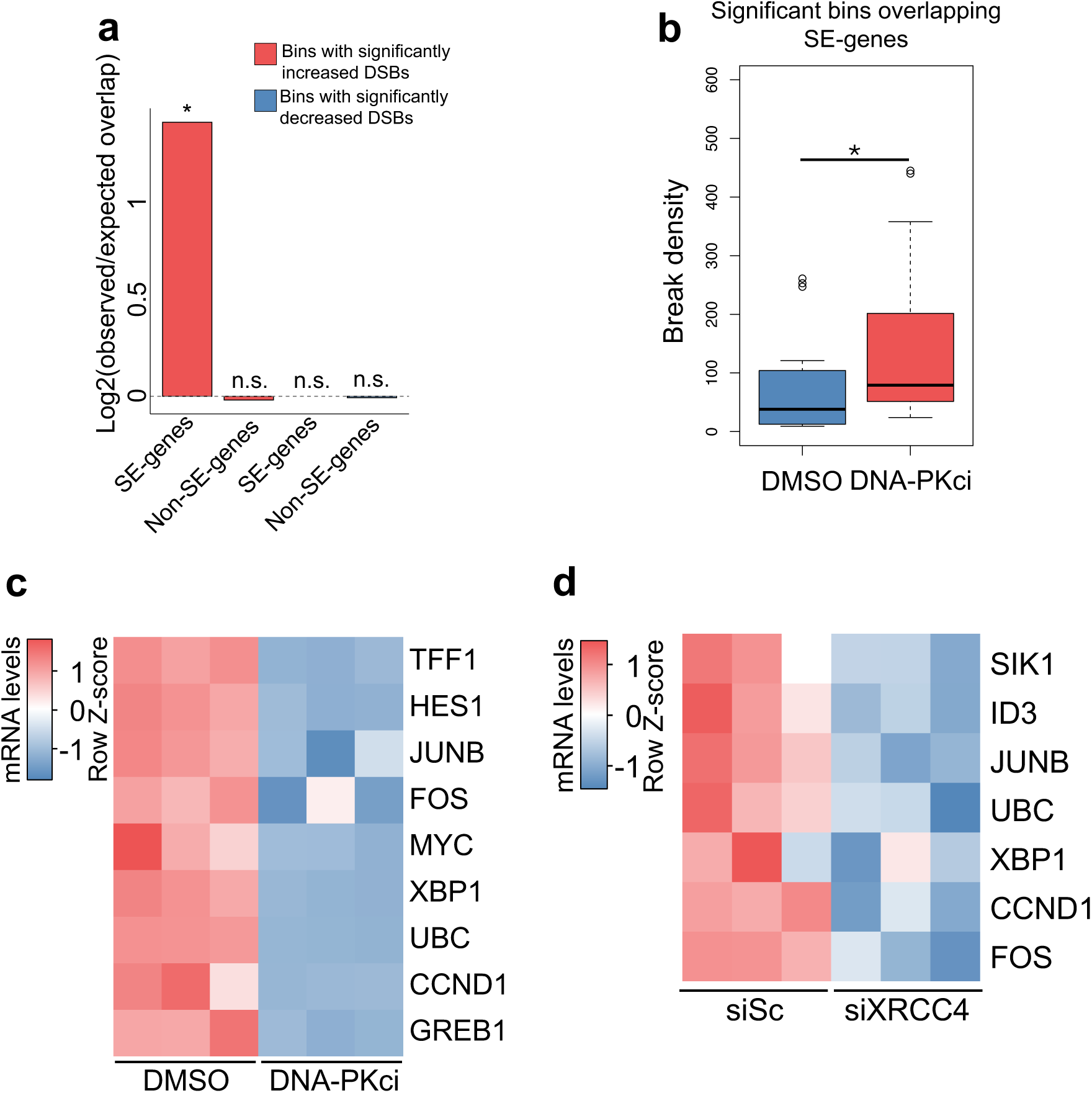
Efficient repair at transcription stress sites is required for SE-regulated oncogene expression. **a, Log2** ratio of observed to expected overlap of significantly affected genomic bins (10KB) with SE-regulated genes and non-SE-regulated genes upon DNA-PKci treatment. P-values were calculated using binomial test. **b**, break density of significantly affected bins overlapping SE-regulated genes. **c**, heatmap showing the change of mRNA levels as measured by spike-in qPCR upon DNA-PKci treatment of top γH2AX-enriched SE-regulated genes in MCF7 cells. d, heatmap showing the change of mRNA levels as measured by qPCR upon XRCC4 KD of top γH2AX-enriched SE-regulated genes in HeLa cells. n.s. indicating p > 0.05, * indicating p < 0.05.

**Extended data Fig. 9.**
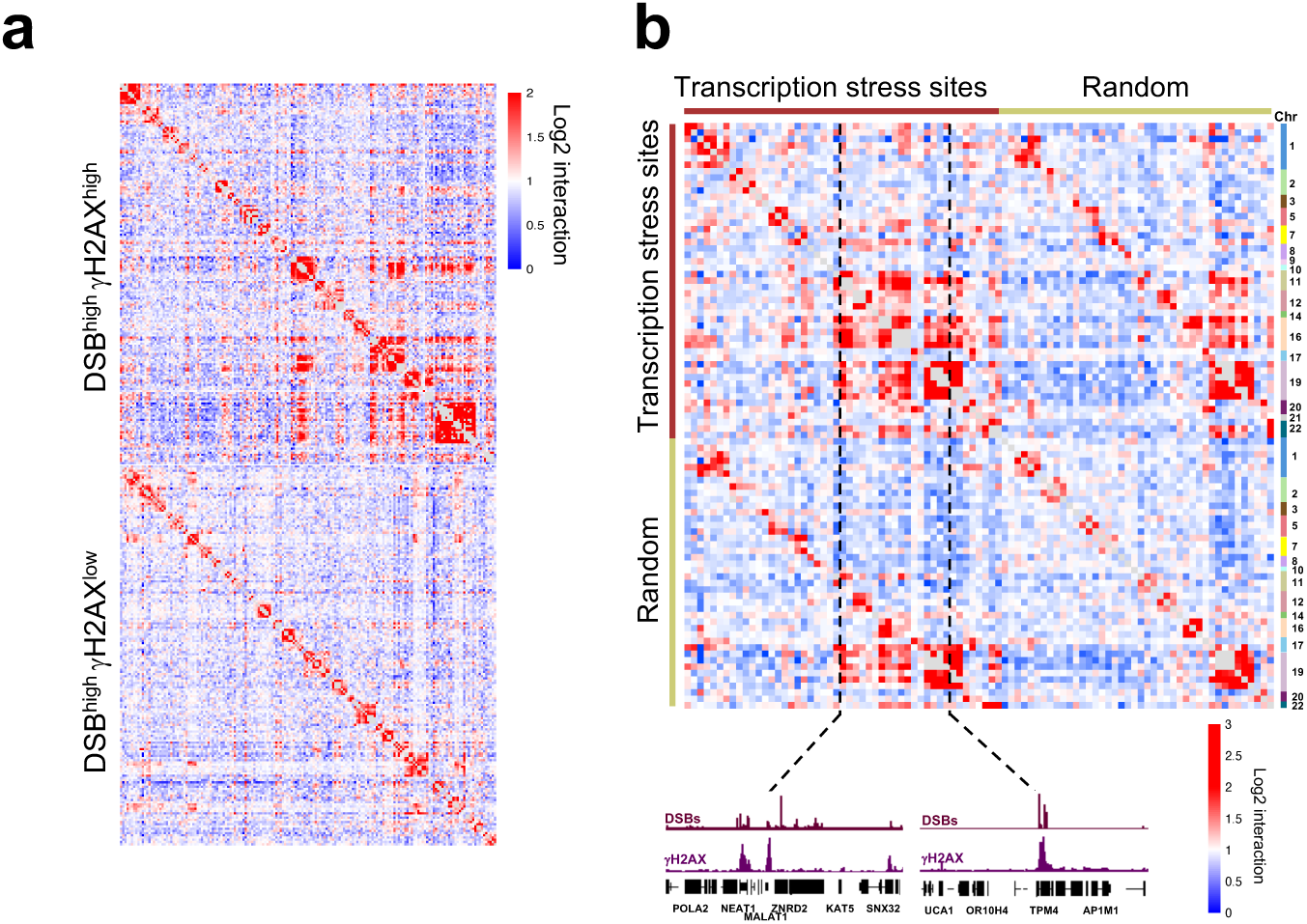
**a**, Heatmap showing Log2 interaction scores for genomic bins containing DSBs^high^ γH2AX^high^ (top) and DSBs^high^ γH2AX^low^. (bottom). **b**, Heatmap showing Log2 interaction scores for genomic bins containing transcription stress sites or randomly selected bind with similar chromosomal distribution.

